# LonP1 Orchestrates UPR^mt^ and UPR^ER^ and Mitochondrial Dynamics to Regulate Heart Function

**DOI:** 10.1101/564492

**Authors:** Bin Lu, Fugen Shangguan, Dawei Huang, Shiwei Gong, Yingchao Shi, Zhiying Song, Lianqun Jia, Juan Xu, Chaojun Yan, Tongke Chen, Mingjie Xu, Yujie Li, Shengnan Han, Nan Song, Pingyi Chen, Lu Wang, Yongzhang Liu, Xingxu Huang, Carolyn K. Suzuki, Zhongzhou Yang, Guanlin Yang

## Abstract

Protein quality control is pivotal to cellular homeostasis and integrity of cardiomyocytes for maintenance of normal heart function. The unfolded protein response (UPR) is an adaptive process to modulate protein quality control in the endoplasmic reticulum (ER) and mitochondria, and is accordingly termed UPR^ER^ and UPR^mt^, respectively. Lon protease (LonP1) is a highly conserved mitochondrial protease to modulate UPR^mt^, which is involved in regulating metabolism, mitophagy, and stress response. However, whether LonP1 regulates UPR^ER^ remains elusive. To investigate the regulation of protein quality control in cardiomyocytes, we generated cardiac-specific LonP1 deletion mice. Our findings show that LonP1 deficiency caused impaired mitochondrial respiratory function and fragmentation. Surprisingly, both UPR^ER^ and UPR^mt^ is substantially induced in LonP1-deletion heart suggesting of LonP1 as a novel regulator of UPR^ER^; however, the activation of UPR^ER^ occurs earlier than UPR^mt^ in response to LonP1 deletion. Consequently, cardiac-specific LonP1 deficiency causes aberrant metabolic reprogramming of cardiomyocytes, pathological heart remodeling, as well as impeded heart function. Thus, we uncovered the novel function of LonP1 as an UPR^mt^ mediator, and reciprocal orchestration of UPR^mt^ and UPR^ER^ and mitochondrial dynamics regulated by LonP1 in the cardiomyocytes that is critical to maintain heart function, which offers exciting new insights into the potential therapeutic strategy for heart failure.

## INTRODUCTION

Mitochondria are important intra-cellular organelles present in most eukaryotic cells, which play a crucial role in ATP production through oxidative phosphorylation (OXPHOS) as well as in determining cell survival or death [1, 2]. Disruption of mitochondrial homeostasis has been implicated in numerous diseases including heart failure. The mitochondrial OXPHOS system produces most of cellular energy (ATP) in almost all cell types. The OXPHOS dysfunction causes a group of multi-systemic diseases which are often progressive or fatal disorders with defects in the cellular ATP supply [3]. Given that the heart is highly dependent on aerobic metabolism, it is particularly susceptible to mitochondrial dysfunction. Energy depletion has been highlighted as an important contributor to the pathology of heart failure [4]. Oxidative stress is believed to be a major causative factor leading to mitochondrial dysfunction. There is accumulating evidence indicating that increased oxidative stress is involved in cardiac hypertrophy and dysfunction [5], and reducing of oxidative stress prevents left ventricular remodeling and dysfunction [5]. There has been considerable interest in the identification of proteins involved and in developing strategies to reduce oxidative stress [6]. The nuclear DNA-encoded mitochondrial ATP-dependent Lon protease (LonP1) belongs to the AAA^+^ family of proteins (ATPase associated with various cellular activities), which exhibits a quality-control protease in selectively eliminating certain abnormal proteins and cellular stress responses in mitochondria [7–9]. Under certain circumstance, LonP1 may also degrade some folded [10] and regulatory proteins [11–14]. LonP1 was upregulated in hypoxia-induced cardiomyocytes and downregulation of LonP1 reduced hypoxia-induced cardiomyocyte apoptosis through decreasing reactive oxygen species (ROS) production [15]. In addition, overexpression of LonP1 enhanced ROS generation and induced apoptosis of cardiomyocytes under normoxic conditions [15]. Moreover, LonP1 may act as a fine-tuning for the function of mitochondrial transcription factor A (TFAM) by precisely degradation of TFAM, which is critical for the replication and transcription of mtDNA [9, 16]. Despite its high conservation throughout evolution, the role of LonP1 *in vivo* in physiological (normal) condition and the pathogenesis of human diseases remains largely unknown. Recent studies reported that LonP1 protein and/or mRNA levels are increased in a variety of tumors [17–21] and LonP1 controls tumor bioenergetics by reprogramming mitochondrial OXPHOS complexes [22]. Furthermore, a link between LonP1 and human diseases has been reported in several genetic diseases [23–29]. Heart failure is closely related to mitochondrial dysfunction; however, the underlying molecular mechanism remains poorly understood. In a mouse model with pressure overload heart failure, the protein carboxylation, tyrosine nitration and cysteine oxidation of LonP1 increased and the protease activity markedly reduced in the isolated mitochondria, which led to mitochondrial respiration deficiency and left ventricular contractile dysfunction [30]. By using haploinsufficiency of LonP1 (LONP1^+/-^) and cardiac-specific overexpression of LonP1 (LonTg) mice model, Venkatesh *et al* found that upregulation of LonP1 alleviates cardiac injury by preventing oxidative damage of proteins and lipids, preserving mitochondrial redox balance and reprogramming bioenergetics by reducing Complex I content and activity [31]. However, the role of LonP1 in the pathogenesis of cardiomyopathy and cardiac dysfunction is still unclear.

The unfolded protein response (UPR) is a protective or an adaptive process to modulate protein quality control in the endoplasmic reticulum (ER) and mitochondria, and is accordingly termed UPR^ER^ and UPR^mt^, respectively. Both UPR^ER^ and UPR^mt^ cause global alteration of transcription networks to increase ER and mitochondrial localized molecular chaperones and proteases to promote the recovery of organellar protein homeostasis (proteostasis) of cell to maintain their function properly [32–34]. However, the UPR^mt^ also promotes a rewiring of cellular metabolism that includes an increase in glycolysis and amino acid catabolism genes with a simultaneous repression of TCA-cycle- and OXPHOS-encoding genes potentially to relieve mitochondrial stress and/or alter cellular metabolism to promote survival [35]. Nevertheless, it may have important clinical implications, for example, mitochondrial changes in response to ER-stress conditions have been described for several human diseases, including type II diabetes and Alzheimer’s disease [36].

In the present study, we developed a mouse model lacking LonP1 in cardiomyocyte to characterize its role in the heart. We demonstrate that both UPR^ER^ and UPR^mt^ are activated in the heart tissue of LonP1 deficient mice. Surprisingly, UPR^ER^ is induced earlier than UPR^mt^. Moreover, our results show that cardiac-specific LonP1 deletion impairs mitochondrial fusion and leads to impaired mitochondrial fragmentation and respiratory function. Finally, our data show that LonP1 deficiency in the heart results in aberrant metabolic reprogramming and pathological heart remodeling and eventually progresses to heart failure. Collectively, these findings identify LonP1 as a novel regulator of cardiomyocyte fate and offers exciting new insights into the potential therapeutic strategy for heart failure.

## RESULTS

### Generation of Cardiomyocyte-Specific LonP1 knockout mice

To investigate the function of LonP1 in the heart, we generated mice deficient in LonP1 specifically in the heart by mating *LonP1*^*LoxP/LoxP*^ mice with cardiomyocyte-specific-MHC-Cre (αMHC-Cre) mice (Fig 1A and 1B). The loss of LonP1 was confirmed at both the transcript and protein levels. Levels of LonP1 mRNA and protein were extremely low and almost undetectable in the heart of LonP1 conditional knockout mice (cLKO) compared with that in littermate controls (*LonP1*^*LoxP/LoxP*^, WT) after two weeks (Fig 1C and 1D). However, we found that there was no difference in LonP1 expression in the other organs of cLKO and WT mice (Fig 1E). Thus, our results indicate that LonP1 was efficiently and specifically deleted in the cardiomyocytes of cLKO mice.

**Fig 1.**
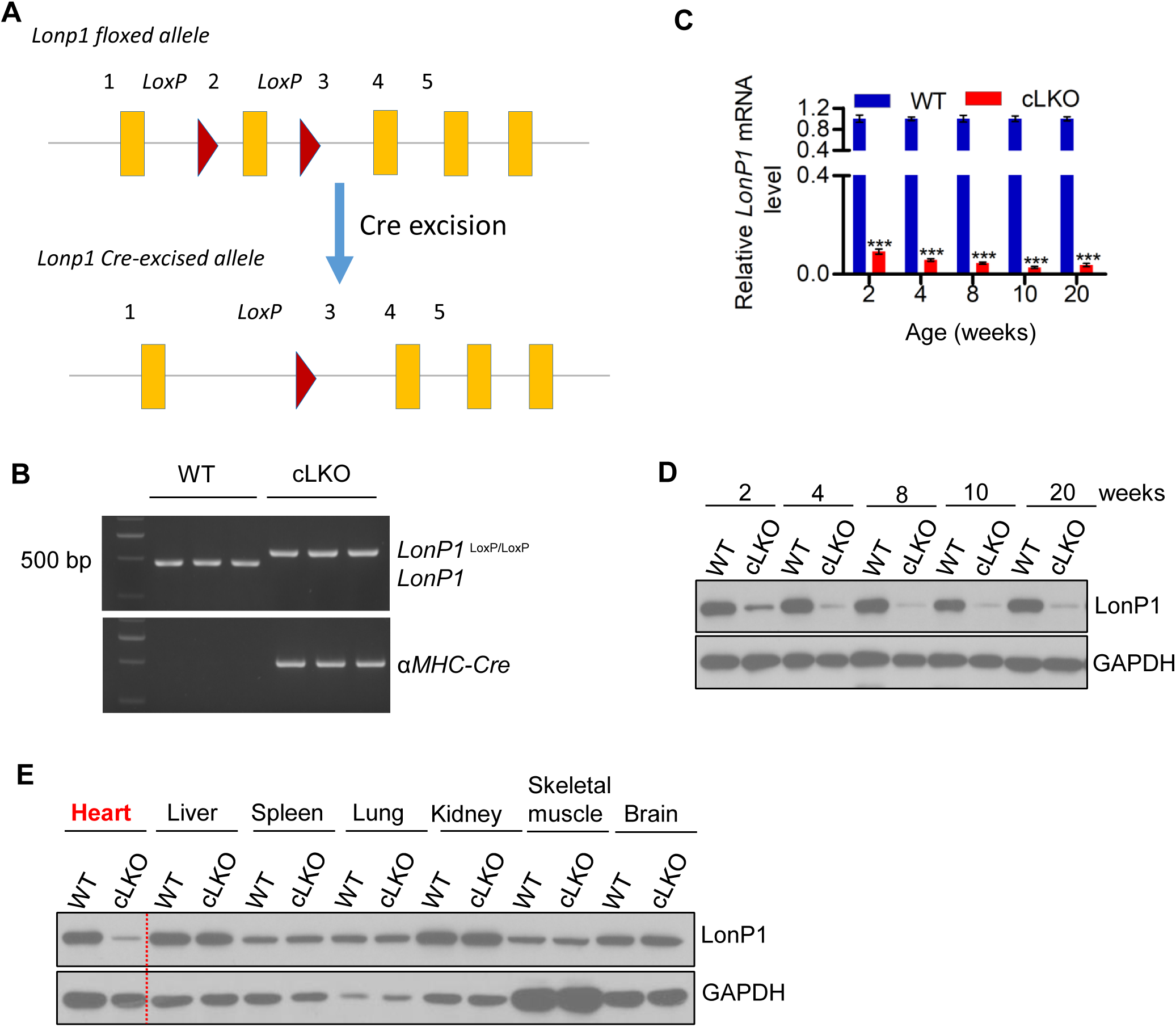
Generation of Cardiomyocyte-Specific LonP1 Deletion Mice. (A) Generation of cardiomyocyte-specific LonP1-deficient mice. *Lonp1*^*LoxP/LoxP*^ mice were crossed with cardiomyocyte-specific-MHC-Cre (αMHC-Cre) mice for deletion of LonP1 in the cardiomyocytes. (B) PCR products were electrophoresed on a 1% agarose gel. Genotype identification shows *Lonp1*^*LoxP/LoxP*^ alleles located at 573 bp, wild type alleles located at 473 bp, and the α-MHC-cre alleles located in 501 bp. *Lonp1*^*LoxP/LoxP*^ with α*-MHC-cre* refers to cLKO mice. (C and D) The loss of LonP1 was confirmed at both the transcript (C) and protein (D) levels. Levels of *Lonp1* mRNA and protein were extremely low and almost undetectable in the hearts of LonP1-conditional knockout mice (cLKO) compared with littermate controls (*Lonp1*^*LoxP/LoxP*^, WT) after two weeks. (E) There was no difference in *LonP1* expression in the other organs between cLKO and WT mice. In (C), data are presented as the means ± SEM (n = 3, \*\*\**p* < 0.001, statistically significant by Student’s t test).

### Cardiomyocyte-Specific Deletion of LonP1 Causes Pathological Heart Remodeling and Impaired Heart Function

We next examined heart function from an early age up to 60 weeks of age in cLKO and WT mice. There were no significant differences observed in the heart weight-to-body weight ratio between WT and cLKO mice before 10 weeks (Fig 2A). However, the cLKO mice at 20 weeks developed cardiomyopathy characterized by increased heart size with age (Figs 2B, S1A and S1B). Based on hematoxylin and eosin (H&E) staining of WT and cLKO mice hearts at different ages, we found a slight increase in the left ventricular (LV) wall thickness in 20-week-old cLKO mice and a significant increase in the cardiac size and LV dilation in 60-week-old cLKO mice (Figs 2C, S1A and S1B). We observed myocardial fibrosis, as shown by Sirius Red and Masson’s trichrome staining, in the hearts of cLKO mice at 20 and 60 weeks (Figs 2D, S1C and S1D).

**Fig 2.**
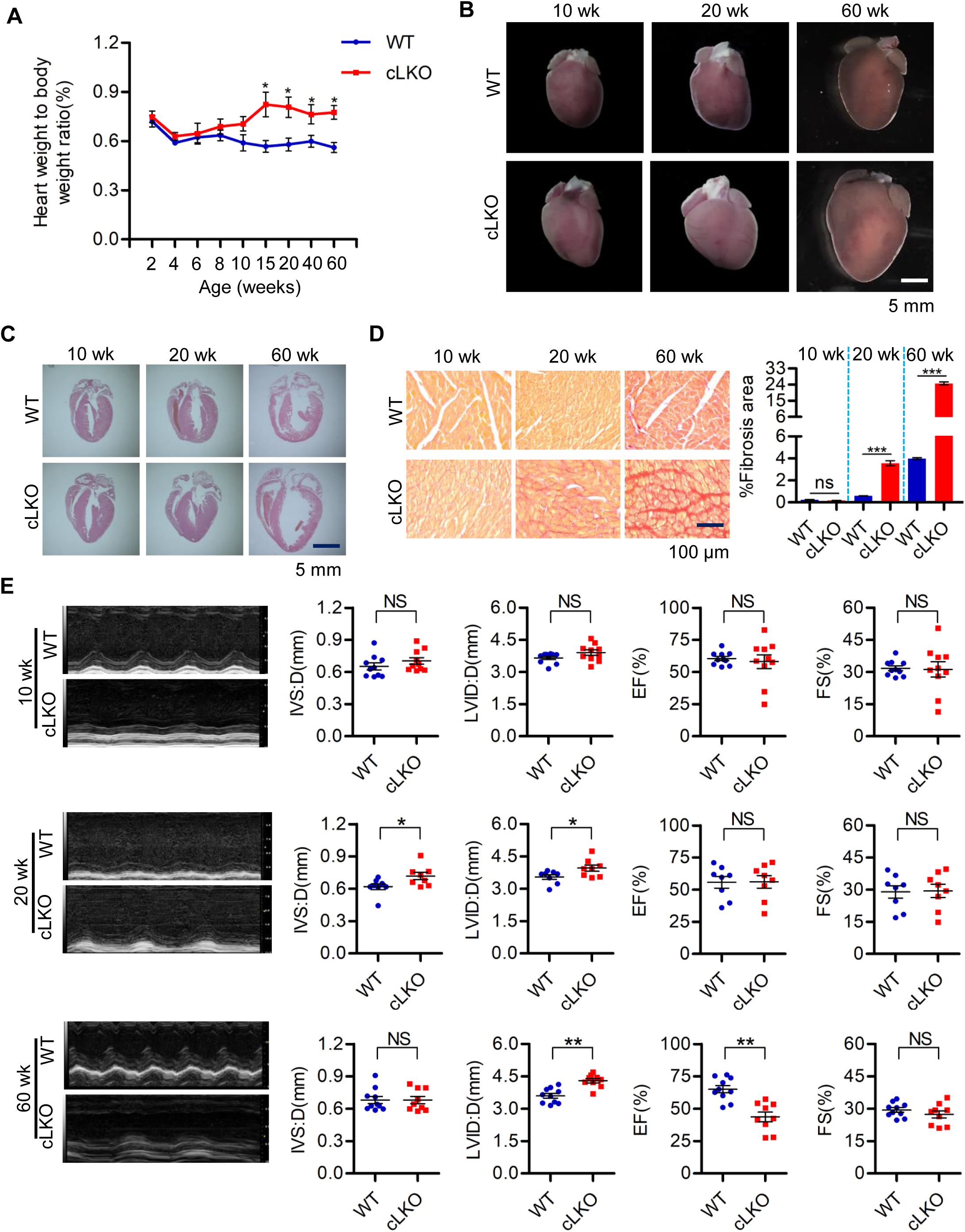
Cardiomyocyte-Specific Deletion of LonP1 Causes Pathological Heart Remodeling and Impaired Heart Function. (A) Heart weight-to-body weight ratios in WT and cLKO mice of 2, 4, 6, 8, 10, 15, 20, 40, and 60 (n=6, \**p* < 0.05, statistically significant by Student’s t test) weeks of age. (B) Heart morphologies of 10-, 20- and 60-week-old WT and cLKO mice. (C) Hematoxylin and eosin staining of WT and cLKO hearts of 10-, 20- and 60-week-old cLKO and WT mice. (D) Representative images showing Sirius Red staining of the cardiac tissues from 10-, 20-, and 60-week-old cLKO mice and their littermate controls. Scale bars, 100 μm. Quantitative analysis of myocardial fibrosis in 10-, 20-, and 60-week-old WT and cLKO mice. Data shown are means ± SEM (n=3, \*\*\**p* < 0.001, statistically significant by Student’s t test). (E) Representative images of transthoracic M-mode echocardiographic tracings and echocardiographic parameters of 10-, 20-, and 60-week-old WT and cLKO mice. IVS:D represents the interventricular septal thickness; LVID:D represents the left ventricular internal diameter in diastole; EF indicates the ejection fraction; and FS indicates the left ventricular fraction shortening. Data shown are means ± SEM (n=8-10, \**p* < 0.05, \*\**p* < 0.01, statistically significant by Student’s t test). See also fig. S4.

Serial echocardiography analyses revealed progressive heart dysfunction in cLKO mice, which became apparent at 20 weeks and was characterized by a statistically significant increase of the interventricular septal depth (IVS:D) and left ventricular internal dimension (LVID), although the left ventricular ejection fraction (EF%) was still normal compared with the WT mice. However, cLKO mice at 60 weeks exhibited severe LV dilation, accompanied by significant EF% reduction (Fig 2E). In conclusion, the cardiomyocyte-specific deletion of LonP1 leads to dilated cardiomyopathy (DCM) and heart failure, which reveals that LonP1 is indispensable for heart function.

### LonP1 Deletion Leads to Mitochondrial Fragmentation

Mitochondrial fission and fusion are critical for mitochondrial function, distribution, and turnover. The balance between fission and fusion ensures the health of cardiac myocyte. Cardiomyocyte specific deletion of fusion proteins Mitofusin 1 and 2 (MFN1 and MFN2), optic atrophy 1 (Opa1), and OPA1-processing peptidases YME1L and OMA1, in adults mice causes mitochondrial fragmentation, and which results in the mice developing dilated cardiomyopathy and heart failure [37–41]. We therefore determined the protein levels of the mitochondrial fission and fusion related proteins dynamin-related protein 1 (DRP1), MFN1 and MFN2, as well as OPA1 and OPA1-processing peptidases OMA1 in cLKO mice by immunoblotting.

To validate the effect of LonP1 ablation to the mitochondrial morphology, we knocked down LonP1 stably in H9c2 rat cardiomyocytes. Using confocal microscopy, we observed significant mitochondrial fragmentation in sh*LonP1* H9c2 cells compared with sh*Cont* H9c2 cells (Figs 3A, 3B and S2A). We further confirmed that the loss of LonP1 in cardiomyocytes of 4- and 10-week-old mice caused mitochondrial fragmentation by using Transmission Electron Microscopy (TEM) (Fig 3C and 3D).

**Fig 3.**
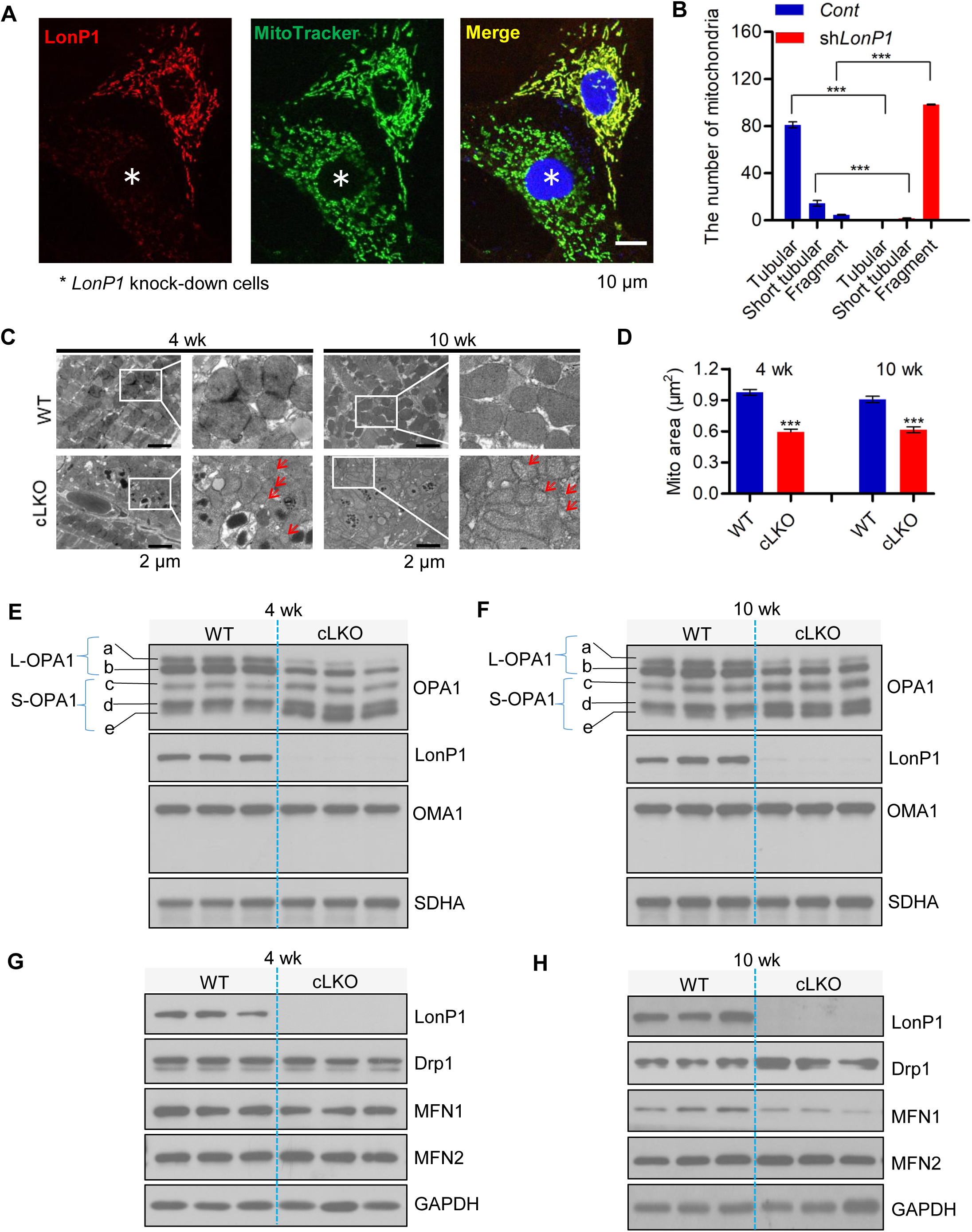
LonP1 Deficiency Leads to Mitochondrial Fragmentation. (A) The LonP1 protein levels and mitochondrial morphologies in control and LonP1 knockdown H9c2 cells were investigated by immunofluorescence (IF) with specific LonP1 antibody and MitoTracker Green. Representative confocal microscopy images are shown (scale bars, 10 μm). Data shown are means ± SEM (n=3, \*\*\**p* < 0.001, statistically significant by Student’s t test). (B) Mitochondrial morphologies described in (A) were counted according to the criteria detailed in the Experimental Procedures. (C) Representative TEM images of 4- and 10-week-old cLKO and WT hearts. Arrows indicate mitochondria with typical morphology. Scale bars, 2 μm. (D) The mitochondrial area of TEM images in (C) was quantified by ImageJ. Data shown are means ± SEM (n=3, \*\*\**p* < 0.001, statistically significant by Student’s t test). (E and F) Western blot analysis of OPA1, OMA1, SDHA and LonP1 protein levels in the heart tissue of 4-(E) and 10-week-old (F) cLKO mice and WT mice. GAPDH was used as a loading control. (G and H) Western blot analysis of Drp1, MFN1, MFN2, and LonP1 protein levels in the heart tissue of 4-(G) and 10-week-old (H) WT and cLKO mice. GAPDH was used as a loading control. See also S2 Fig.

Furthermore, we observed a significant reduction in L-OPA forms a and b, and S-OPA forms c and e were increased dramatically, whereas S-OPA1 form d remained unchanged in the hearts of 4- and 10-week-old cLKO mice in comparison to control mice. OMA1 is a protease that cleaves OPA1 from L-OPA1 to S-OPA1 [42]; however, no changes were observed in OMA1 expression in the hearts of 4- and 10-week-old cLKO mice (Fig 3E and 3F). Next, we examined Drp1, a protein that induces mitochondrial fission, as well as MFN1 and MFN2, which promote mitochondrial fusion in the hearts of 4- and 10-week-old cLKO and control mice. We found that there was no difference in protein expression of Drp1, MFN1, and MFN2 in the hearts of 4-week-old cLKO and control mice, whereas the protein expression of Drp1 was statistically significantly increased, and MFN1 was statistically significantly decreased in the hearts of 10-week-old cLKO mice compared with control mice, while there were no differences in MFN2 (Fig 3G and 3H). Notably, there was no difference in any of these proteins involved in mitochondrial fusion and fission in the hearts of 2-week-old cLKO mice and their littermate controls (S2B and S2C Fig).

These results together indicate that LonP1 ablation in cardiomyocytes promotes OPA1 processing and Drp1 expression, as well as reduces MFN1 expression to enhance mitochondrial fragmentation.

### Loss of LonP1 Induces UPR^ER^ Prior to UPR^mt^

To further understand the effects of LonP1 deletion to mitochondria and ER, we subsequently examined the UPR^ER^ and UPR^mt^ at different postnatal stages of mice. To test this, we determined the expression level of UPR^ER^-related proteins in cLKO mice and controls. We observed that inositol-requiring enzyme 1α (IRE1α), eukaryotic initiation factor 2α subunit (eIF2α), and the phosphorylation of eIF2α (p-eIF2α), activating transcription factors 4 and 6 (ATF4 and ATF6), increased significantly in response to LonP1 depletion in 4- and 10-week-old cLKO mice (Fig 4A and 4B). In addition, we observed that protein disulfide isomerase (PDI) remained unchanged in 4-week-old cLKO and control mice, whereas the PDI protein level was significantly increased in 10-week-old cLKO mice (Figures 4A and B).

**Fig 4.**
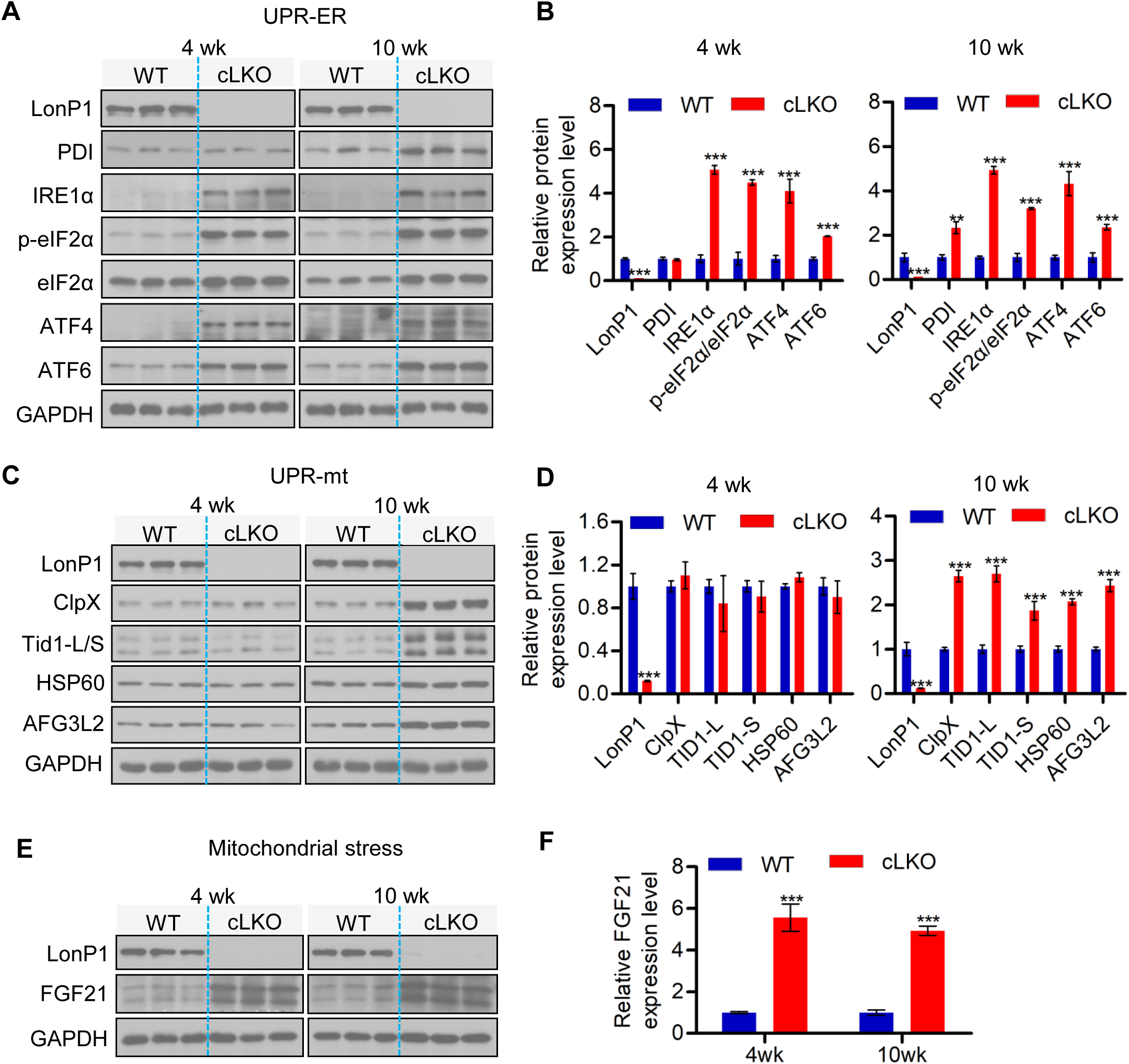
Deletion of LonP1 Induces UPR^ER^ and UPR^mt^. (A and B) Western blot analysis (A) and quantification (B) of endoplasmic reticulum (ER) stress proteins PDI, IRE1α, p-eIF2α, ATF4 and ATF6 in WT and cLKO hearts at 4 and 10 weeks of age. GAPDH was used as a loading control. Data shown are means ± SEM (\*\**p* < 0.01, \*\*\**p* < 0.001, statistically significant by Student’s t test). (C and D) Western blot analysis (C) and quantification (D) of LonP1 and the mitochondrial unfolded protein response related proteins ClpX, Tid1-L/S, HSP60, and AFG3L2 in WT and cLKO hearts at 4 and 10 weeks of age. GAPDH was used as a loading control. Data shown are means ± SEM (\*\*\**p* < 0.001, statistically significant by Student’s t test). (E and F) Western blot analysis and quantification of the mitochondrial stress marker FGF21 protein level in WT and cLKO hearts at 4 and 10 weeks of age. GAPDH was used as a loading control. Data shown are means ± SEM (\*\*\**p* < 0.001, statistically significant by Student’s t test). See also S3 Fig.

We next examined the UPR^mt^-related proteins in cLKO and control mice. We found that ClpX, Tid1-L/S, HSP60, and the ATPase family gene 3-like 2 (AFG3L2) protease involved in the turnover of unfolded proteins, were significantly increased in 10-week-old cLKO mice. Surprisingly, the expression levels of these UPR^mt^-related proteins remained unchanged in 4-week-old cLKO mice (Fig 4C and 4D). Notably, we observed that UPR^ER^- and UPR^mt^-related protein levels remained unchanged between 2-week-old cLKO mice and their littermate controls (Figures S3A and S3B). These results suggest that the cardiomyocyte-specific loss of LonP1 activates both UPR^ER^ and UPR^mt^; however, the activation of UPR^ER^ is more sensitive than that of UPR^mt^ to the loss of LonP1 in cardiomyocytes.

In addition, we evaluated the expression level of fibroblast growth factor 21 (FGF21), a marker of mitochondrial stress signal [43, 44], in 4- and 10-week-old WT and cLKO mice, and we observed a strong increase of FGF21 protein levels in the hearts of 4- and 10-week-old cLKO mice (Fig 4E and 4F). Similar to the case for UPR^ER^- and UPR^mt^-related proteins, there was no difference in FGF21 protein levels between 2-week-old cLKO mice and their littermate controls (S3C Fig).

Taken together, our results reveal that LonP1 ablation in cardiomyocytes activates both UPR^ER^ and UPR^mt^; however, LonP1-deletion induces UPR^ER^ prior to UPR^mt^. Although LonP1 located predominantly in the mitochondrial matrix, it may play a potential pivotal role in maintaining the homeostasis of the ER and mitochondria. Our findings also indicate that cytokine FGF21 is, indeed, an early marker of mitochondrial dysfunction, suggesting that FGF21 could be a direct target of UPR^mt^.

### Cardiomyocyte-specific deletion of LonP1 leads to abnormal mitochondrial morphology and dysfunction

We proceeded to investigate the mitochondrial morphology and respiratory chain function in the mitochondria of the hearts of cLKO mice. Similarly, TEM analysis showed increased fragment mitochondria in the LonP1-deletion hearts, which indicated abnormal mitochondrial dynamics (Fig 5A). Moreover, we observed distorted mitochondrial morphology in cLKO mice hearts (Fig 5A). We further observed extensive electron-dense aggregates within the mitochondria of cLKO mice hearts but not in other cell organelles, and no inclusions were observed in WT mice (Fig 5A). In the heart mitochondria of 20-week old cLKO mice, the aggregates were significantly increased. Moreover, we found that some large mitochondria were scattered, with dense deposits of different sizes filling the mitochondrial matrix, and the mitochondrial ridges significantly decreased or even disappeared in the hearts of 60-week-old cLKO mice (Fig 5A).

**Fig 5.**
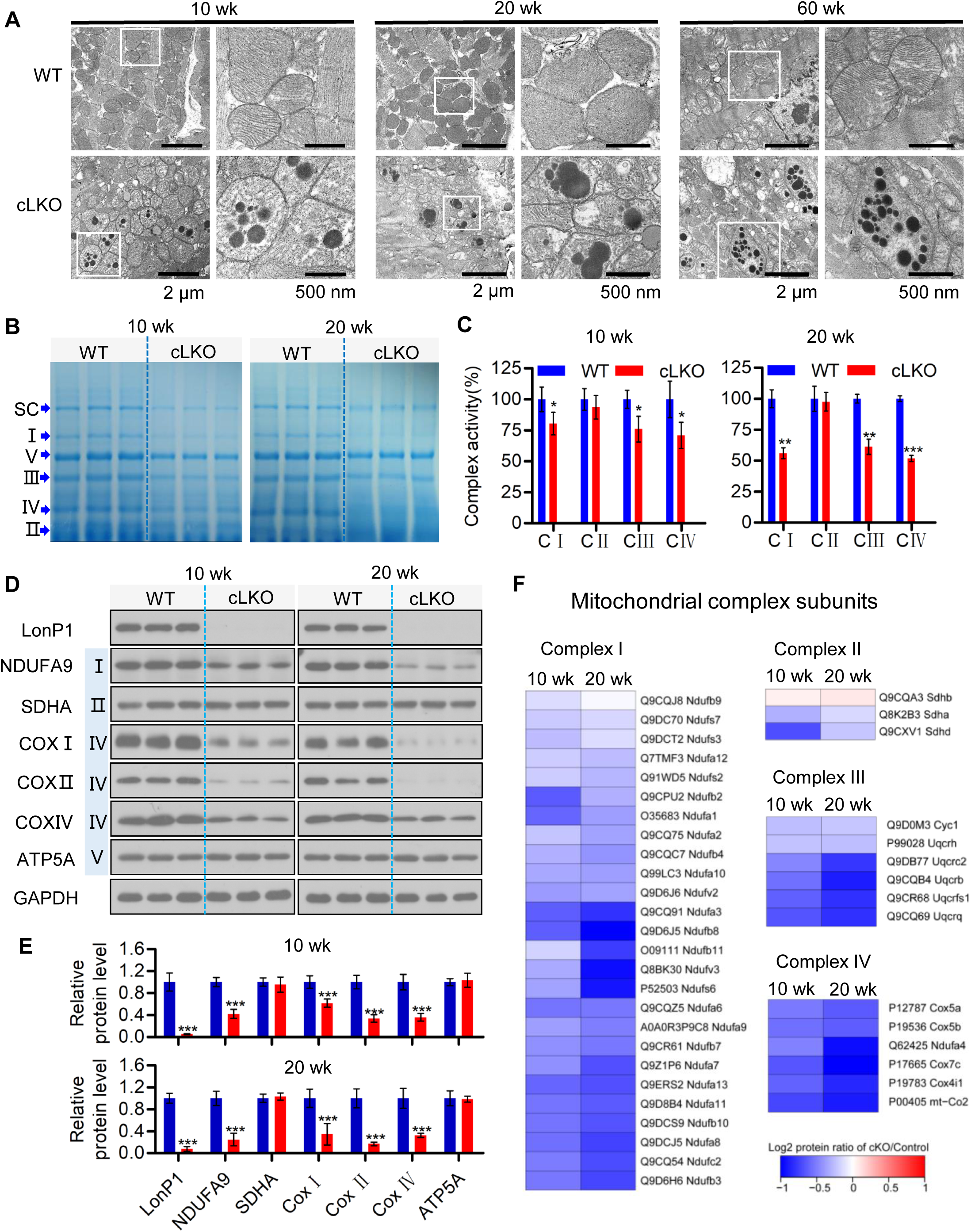
Cardiomyocyte-Specific Deletion of LonP1 Leads to Abnormal Mitochondrial Morphology and Dysfunction. (A) TEM of 10-, 20-, and 60-week-old WT and cLKO hearts. Scale bars, 2 μm and 500 nm, respectively. (B) Coomassie Brilliant Blue staining of BN-PAGE with indicated positions and quantifications of specific complexes I to V of WT and cLKO hearts at 10 and 20 weeks of age. (C) Enzymatic activity of complexes I to IV of the mitochondrial electron transport chain (ETC) from the heart tissue of 10- and 20-week-old WT and cLKO mice. Data shown are means ± SEM (n=3, \**p* < 0.05, \*\**p* < 0.01, \*\*\**p* < 0.001, statistically significant by Student’s t test). (D and E) Western blot analysis (D) and quantification by ImageJ (E) of protein levels for representative ETC complex subunits from the heart tissue of 10- and 20-week-old WT and cLKO mice. GAPDH was used as the loading control. Data shown are means ± SEM (n=3, \*\*\**p* < 0.001, statistically significant by Student’s t test). (F) Heatmap showing protein abundance changes in the subunits of complexes I (C I), II (C II), III (CIII), and IV (C IV) obtained via high-throughput quantitative proteomics of enriched mitochondrial preparations. The relative abundance changes in WT and cLKO mice are presented using the protein ratio in relation to each control at 10 and 20 weeks of age.

We next examined the steady-state level of assembled OXPHOS complexes by blue native PAGE (BN-PAGE) analysis and subsequent examination of mitochondrial respiratory chain (MRC) enzyme activities. We observed a significant decrease of the super complex (SC) and complex I, III, and IV levels in the heart tissue of both 10- and 20-week-old cLKO mice (Fig 5B); however, the complex II and V levels only decreased slightly in the heart tissue of both 10- and 20-week old cLKO mice (Fig 5B). The enzymatic activities of heart tissue mitochondrial complex I, III, and IV decreased dramatically in 10-week-old cLKO mice and which was further decreased in 20-week-old cLKO mice (Fig 5C).

We also detected the protein expression level of several MRC enzyme subunits by Western blotting and carried out quantitative analysis. Consistent with the results of BN-PAGE and MRC enzyme activity analyses, we found that SDHA and ATP5a encoded by the nuclear gene remain unchanged, and the mtDNA encoded subunit COX I and COX II, as well as the nuclear gene-encoded subunit NDUFA9 and COX IV, were significantly decreased, with the downregulation in 20-week-old more apparent than that in 10-week-old mice (Fig 5D and 5E). Finally, we performed quantitative proteomic analysis of the heart tissue of LonP1-knockout mice using isobaric tags for relative and absolute quantization (iTRAQ) and observed that LonP1 knockout reduced most of the mitochondrial complex I, II, III, and IV subunit expression levels (Fig 5F). Moreover, these mitochondrial complex subunits were further reduced in 20-week-old cLKO mice, except for the three complex II subunits detected by iTRAQ (Fig 5F).

These results together demonstrate that LonP1 knockout in the myocardium induces the accumulation of protein aggregates, impairing mitochondrial structure and morphology, as well as MRC function.

### Loss of LonP1 in Cardiomyocytes Leads to Metabolic Reprogramming through Enhancing Glycogenesis and Amino Acid Metabolism

Our data showed that LonP1 depletion suppressed H9c2 cell proliferation (Fig 6A and 6B), promoted both intracellular ROS and mitochondrial superoxide generation and caused mitochondrial depolarization (Fig 6C, 6D and 6E). These results strongly indicate that LonP1 is essential for mitochondrial homeostasis and that the loss of LonP1 may cause oxidative damage in H9c2 cells. Mitochondria are the major source of ATP generation, and to further evaluate the impacts of LonP1 on mitochondrial function, we analyzed cellular oxygen consumption rates (OCR) in control and LonP1-knockdown H9c2 cells. We detected that silencing LonP1 dramatically decreased the overall OCR. We further assessed the various parameters of mitochondrial function by analyzing OCR data at each time point. Our results showed that basal respiration, maximum respiration and ATP production were markedly decreased (Fig 6F and 6G) in LonP1-depleted H9c2 cells.

**Fig 6.**
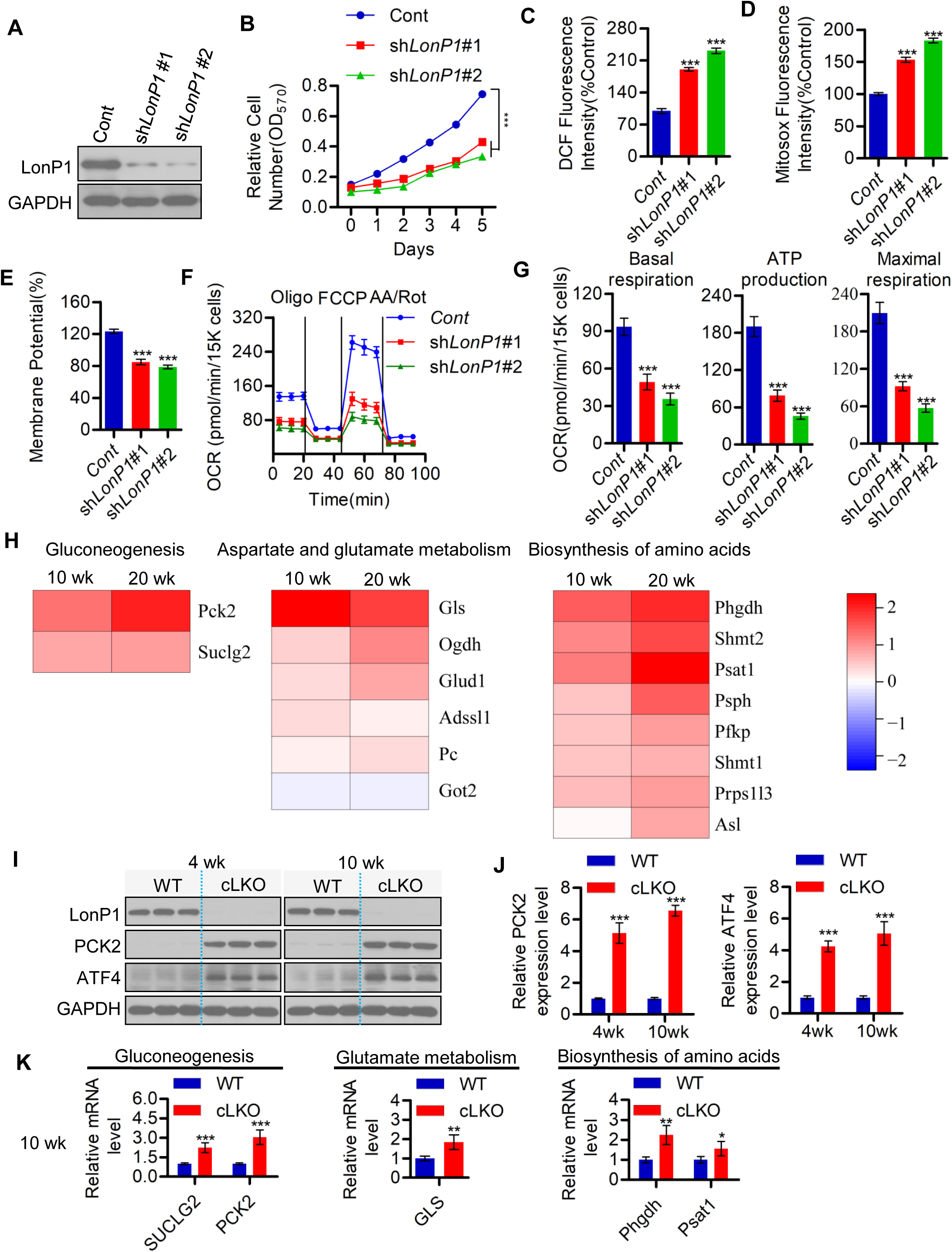
Loss of LonP1 in Cardiomyocytes Leads to Metabolic Reprogramming through Enhancing Glycogenesis and Amino Acid Metabolism. (A) Western blot analysis demonstrating LonP1 stable knockdown in H9c2 cells. (B) Cell proliferation of stable LonP1-knockdown H9c2 cells determined using an MTT Cell Proliferation and Cytotoxicity Assay Kit according to the manufacturer's instructions. Data shown are means ± SEM (n=3, \*\*\**p* < 0.001, statistically significant by Student’s t test). (C) LonP1 stable knockdown increases intracellular ROS (H_2_O_2)_ production in H9c2 cells, as measured using Reactive Oxygen Species Assay Kit (DCFH-DA). Data are plotted as percentages of increase in the median fluorescence intensity (MFI) and are shown as the means ± SEM (n = 3, \**p* < 0.05, \*\**p* < 0.01). (D) Mitochondrial superoxide levels of control and Lon knockdown stable H9c2 cells were detected by MitoSox staining and analyzed by flow cytometry. Data are plotted as percentages of alteration of the mean fluorescence intensity (MFI) and are shown as the means ± SEM (n = 3, \**p* < 0.05, \*\**p* < 0.01). (E) LonP1 stable knockdown reduces the mitochondrial membrane potential (MMP) of H9c2 cells. Cells were stained with JC-1 and analyzed by flow cytometry. The ratio of fluorescence intensities Ex/Em = 490/590 and 490/530 nm (FL590/FL530) were recorded to show the MMP level of each sample. Data are presented as the means ± SEM (n = 3, \*\*\**p* < 0.001). (F and G) The intact cellular oxygen consumption rate (OCR) of H9c2 cells in the indicated conditions were measured in real time using the Seahorse XF96 Extracellular Flux Analyzer. Basal OCRs were measured at three time points, followed by sequential injection of the ATP synthase inhibitor oligomycin (1 μM), the uncoupler FCCP (1 μM), the complex I inhibitor rotenone (1 μM) and the complex III inhibitor antimycin A (1 μM). Data shown are means ± SEM (\*\*\**p* < 0.001, statistically significant by Student’s t test). (H) Heatmap showing protein abundance changes in some proteins involved in gluconeogenesis, aspartate and glutamate metabolism and the biosynthesis of amino acids obtained via high-throughput quantitative proteomics of enriched heart tissues. The relative abundance changes in WT and cLKO mice are expressed using the protein ratio in relation to each control at 10 and 20 weeks of age. (I and J) Western blot and quantification of representative protein levels of ATF4 and PCK2 in WT and cLKO hearts at 4 and 10 weeks, using GAPDH as the loading control. Data shown are means ± SEM (n = 3, \*\*\**p* < 0.001, statistically significant by Student’s t test). (K) qRT-PCR analysis the transcripts of SUCLG2, PCK2, GLS, PHGDH, and PSAT1 in WT and cLKO hearts at 10 weeks. Data shown are means ± SEM (n = 3, \**p* < 0.05, \*\**p* < 0.01, \*\*\**p* < 0.001, statistically significant by Student’s t test).

To explore the underlying molecular mechanism of the LonP1-driven H9c2 cell phenotype, we performed a quantitative proteomic analysis of heart tissue from WT and LonP1 KO mice using iTRAQ labeling and subsequent LC-MS/MS analysis. iTRAQ data showed significant upregulation of crucial metabolic enzymes, especially those participating in the processes of gluconeogenesis and aspartate and glutamate metabolism, as well as the biosynthesis of amino acids in both 10-week-old and 20-week-old cLKO mice (Fig 6H), indicating that cardiomyocyte-specific deletion of LonP1 leads to metabolic reprogramming through enhancing glycogenesis and amino acid metabolism to promote cell survival under excessive oxidative damage stress caused by LonP1 ablation. We further confirmed that the protein levels of mitochondrial phosphoenolpyruvate (PEP) carboxykinase (PEPCK-M or PCK2) and activating transcription factor 4 (ATF4) were increased by Western blotting analysis (Fig 6I and 6J). It was well elucidated that PCK2, a key enzyme in glycogenesis, could be transcriptionally regulated by ATF4 through binding to a putative ATF/CRE composite site within the PCK2 promoter functioning as an amino acid response element, which mediates PCK2 transcriptional upregulation [45]. Consistently, we found that the mRNA transcripts of *SUCLG2*, *PCK2*, *GLS*, *PHGDH* and *PSAT1* were all upregulated in cLKO hearts (Fig 6K). Collectively, our data indicate that the cardiomyocyte-specific deletion of LonP1 promotes metabolic reprogramming to overcome LonP1 deficiency caused mitochondrial dysfunction, excessive oxidative stress, and intermediate metabolites limitations, thus promoting cLKO mouse survival.

## DISCUSSION

Mitochondrial proteins are continuously damaged by exogenous and/or endogenous various stresses, such as ROS, heat, and toxic compounds, which cause proteins damaged due to misfolding or aggregation. Mitochondrial protein quality control system can sense the damaged protein which are selectively degraded by different proteolytic systems within mitochondrial, therefore maintains the protein homeostasis. In the mitochondrial matrix, there are three proteases are responsible for the degradation of damaged mitochondrial proteins, including ClpXP, m-AAA and LonP1, which are members of the ATPase associated with diverse cellular activities (AAA+) superfamily. However, the defects in mitochondrial protein quality control system will lead to accumulation of damaged proteins in the mitochondria, thus causing mitochondrial dysfunction and ultimately leading to cellular dysfunction. In the recent years, LonP1 has been emerged as an essential and fundamental component of mitochondrial quality control system and as an important stress sensor in the mitochondrial alterations that could be observed in a number of chronic diseases and human cancers [8, 19, 46, 47]. A recent *in vitro* study shows that LonP1 is also required for maturation of a subset of mitochondrial matrix proteins and that LonP1 reduction elicits an integrated stress response [48].

In the present study, we demonstrate that cardiomyocyte-specific ablation of LonP1 causes the mice develop DCM and ultimately progresses to heart failure, which indicates the essential roles of LonP1 in normal myocardial cells. Our study shows that LonP1 mediates UPR^ER^ orchestrating UPR^mt^ to regulate heart function and remodeling. The deletion of Lon1 in cardiomyocytes of mice causes imbalance in mitochondrial dynamics, and OXPHOS dysfunction, thus leading to DCM and eventually to heart failure. These findings describe a mechanism by which protein quality control system dysfunction predisposes the myocardium to injury via altered mitochondrial fusion and fission balance as well as susceptibility to chronic mitochondrial stress.

ClpP plays a critical role in the activation of the UPR^mt^ in *Caenorhabditis elegans* [49]. It was also found that ClpX, which multimerizes with ClpP to form the functional ATP-dependent protease ClpXP, could stimulate the UPR^mt^ in mammalian cells similar to the case for the UPR^mt^ in *Caenorhabditis elegans* [50]. A recent report showed that ClpP might be dispensable for mammalian UPR^mt^ initiation since the absence of ClpP could trigger compensatory responses in mice [51]. However, the role of ClpP in mammalian UPR^mt^ is still unclear. LonP1 is also an ATP-dependent mitochondrial matrix protease mainly involved in the degradation of misfolded and aggregated abnormal proteins within the matrix. Here, we show that LonP1 ablation causes many compensatory responses in cLKO mice, including activation of the UPR^ER^. Strikingly, the UPR^ER^ is activated prior to the UPR^mt^, suggesting that LonP1 expression is a fine-tuning sensor of the UPR^ER^ and UPR^mt^. Upregulation of both the UPR^ER^ and UPR^mt^ in cardiomyocytes of LonP1-deletion mice results in well-tolerating mice with no major changes of longevity under unstressed conditions. In addition to the activation of both the UPR^ER^ and UPR^mt^ in the hearts of cLKO mice, FGF21, a key regulator of glucose and lipid metabolism and energy balance, is also increased dramatically to compensate for the deletion of LonP1 expression. The increased expression of FGF21 further ensures cLKO mice survival under unstressed conditions.

Balanced mitochondrial dynamics are critical for normal myocardial function [52]. Mitochondrial fission ensures biogenesis and efficiently removes damaged or old mitochondria through mitophagy, whereas mitochondrial fusion facilitates the mixing and exchange of vital metabolites and mtDNA between different mitochondria to prevent the accumulation of mitochondrial damage in cells, thus properly maintaining the mitochondrial functions [53, 54]. OPA1, located in the inner mitochondrial membrane (IMM) of mitochondria, plays an important role in the regulation of mitochondrial fusion and fission [55]. OPA1 is cleaved by mitochondrial processing peptidase to form the mature isoform of OPA1 (L-isoform) [56], which is further processed into two shorter isoforms, S1 and S2 OPA1. L-OPA1 is sufficient for mitochondrial fusion; however, this feature is lost after cleavage to the S-isoform [56]. LonP1-deletion significantly reduced L-OPA1 through the cleavage of OPA1 and increased L-OPA1 levels, promoting mitochondrial fission. Moreover, Drp1, which is the master regulator of mitochondrial division in most eukaryotic organisms [55], was increased in cardiomyocyte-specific LonP1-deletion mice. These results suggest that the loss of LonP1 triggers stress-induced OPA1 processing and leads to the accumulation of S-OPA1 forms c and e, causing unbalanced fusion and fission of mitochondria and impairs mitochondria morphology in cardiomyocytes lacking LonP1.

We demonstrate that the loss of LonP1 causes severe impairment of mitochondrial function and morphology, promoting metabolic reprogramming to meet energy limitations and intermetabolite supplies. Mitochondrial homeostasis is essential for normal cell physiological equilibrium, and its disruption promotes mitophagy, which contributes to multiple diseases [57]. Our results clearly show that LonP1 ablation in myocardial cells promotes gluconeogenesis and aspartate and glutamate metabolism, as well as the biosynthesis of amino acids, via upregulating multiple vital metabolic catalytic enzymes, such as SUCLG2, GLS, PHGDH, and PAST1. Importantly, mitochondrial PEP-carboxykinase (PCK2), a key enzyme of gluconeogenesis, is dramatically transcriptionally upregulated due to LonP1 depletion, suggesting that LonP1 ablation may drive PCK2-mediated metabolic adaptation in cardiac myocytes to support TCA cycle metabolism and glycolytic intermediates for biosynthesis [58]. The activity of PCK2 depends on mtGTP generated by the SUCLG2 form of SCS in the mitochondrial matrix [59]. We show that the SUCLG2 protein and mRNA expressions are stimulated by LonP1 ablation, indicating a synergistic effect to strengthen PCK2 activity in LonP1-deficient cardiomyocytes. Moreover, we found that ATF4 was upregulated, indicating that PCK2 could be transcriptionally regulated by ATF4 through regulating a putative ATF/CRE composite site containing the PCK2 promoter, which caused transcriptional upregulation of PCK2 [45, 60]. Our results argue that ATF4-induced PCK2 expression could promote LonP1-deletion cardiomyocytes survival in energy- and intermediate metabolites-limited environments. These results suggest that LonP1-deletion cardiomyocytes may activate ATF4/PCK2-mediated metabolic reprogramming to orchestrate energy and intermediate metabolite homeostasis, which contributes to myocardial cell survival.

In the past, many puzzling results have been reported concerning ER stress induced by mitochondrial dysfunction/stress. Frequently, researchers reveal the mitochondrial structural or functional defects induced by ER stress without a reasonable explanation. Here, we unveiled that LonP1-mediated orchestration of UPR^mt^, UPR^ER^ and mitochondrial dynamics, and this finding may provide a clue to this long-standing obscurity (Fig 7).

**Fig 7.**
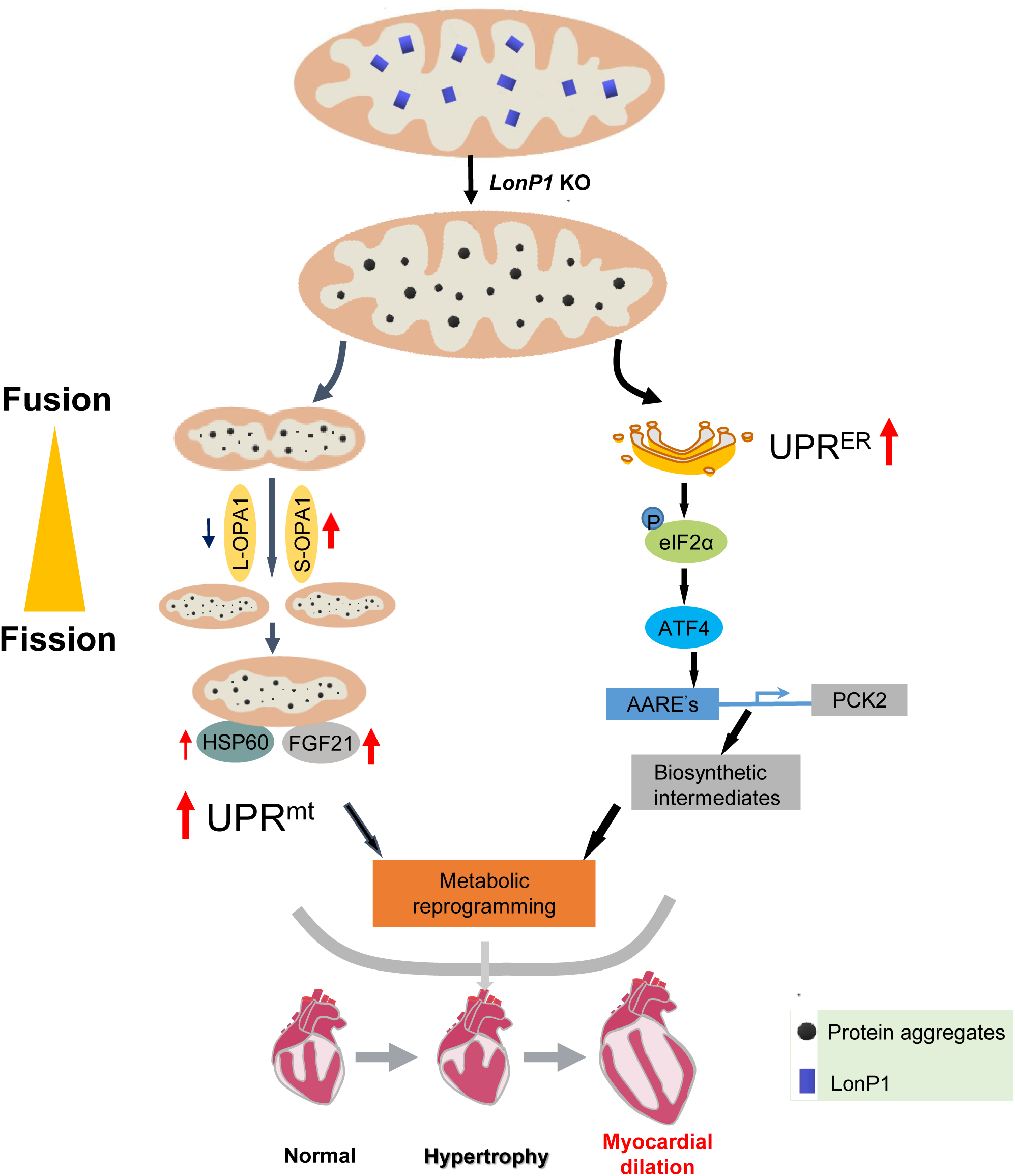
Proposed Model of LonP1 in Modulating Heart Function.

As reduction of LonP1 levels in cardiac myocytes led to DCM and heart failure, it is possible that increased expression of LonP1 may have a cardioprotective effect against DCM and preventing heart failure. It will also be interesting to examine whether there is a correlation between the cardiac level of LonP1 and the incidence of DCM in humans.

## Materials and Methods

### Mice

All mice used in this study were housed at the Wenzhou Medical University Animal Center in a specific pathogen free (SPF) facility with individually ventilated cages. The room in the animal facility has controlled constant temperature (20-25°C) and humidity (30%–70%), with a 12-hour light/dark cycle. Mice were provided ad libitum access to water and normal rodent chow diet. All procedures were reviewed and approved by the Institutional Animal Research Committee of Wenzhou Medical University.

### Construct Method of *Lonp1*^*LoxP/LoxP*^ Mice and Genotyping

The strain was generated as follows. An L1L2_Bact_P cassette coding FRT-lacZ-loxP-neomycin-FRT-loxP was inserted between exon 1 and exon 2 and another loxP was inserted immediately after exon 2. The linearized construct was transfected into C57BL/6J embryonic stem cells and neomycin-resistant clones were screened by PCR. Clones that had undergone homologous recombination were injected into albino C57BL/6J blastocysts and the resulting chimeric mice were crossed with Gt (ROSA) 26Sortm1 (FLP1) Dym (also known as ROSA26::FLPe knockin) mice to excise the neomycin resistance cassette. The PCR primers for genotyping are listed in Table S1.

### Generation of Heart-Specific *Lonp1*-Knockout Mice and Genotyping

The cLKO (*LonP1*^*LoxP/LoxP*^/α*-MHC-cre*) mice were generated by crossing *LonP1*^*LoxP*/+^/α*-MHC-cre* mice with *LonP1*^LoxP/+^ mice. Littermate mice were considered as control mice, except the cLKO mice. The mice were maintained on a mixed C57BL/6.SV129 background. No differences between the characteristics of wild-type (WT) *LonP1*^+/+^ and *LonP1*^LoxP/LoxP^ mice were observed from birth until more than 1 year old. Genomic DNA was extracted from tail snips using a QIAGEN Gentra Puregene Tissue kit, and PCR was performed using the genomic DNA obtained as the template. The PCR primers for genotyping are listed in Table S1.

### Reagents and Antibodies

Glucose was obtained from Sigma. The intact cellular oxygen consumption rate (OCR) assay kit was purchased from Seahorse Bioscience. Horseradish peroxidase (HRP)-conjugated, anti-rabbit, and anti-mouse immunoglobulin G, the ROS assay kit (DCFH-DA), 5,5’,6,6’-tetrachloro-1,1’,3,3’-tetraethylbenzimidazolylcarbocyanine iodide (JC-1), and the MTT Cell Proliferation and Cytotoxicity Assay kit were obtained from Beyotime. The BCA Protein Assay Kit and Pierce ECL Western Blotting Substrate were obtained from Thermo Fisher Scientific. Primary antibodies were obtained from the following suppliers: anti-NDUFA9 (20312-1-AP, Proteintech); anti-Cox II (55070-1-AP, Proteintech); anti-ATP5A (14676-1-AP, Proteintech); anti-MFN1 (13798-1-AP, Proteintech); anti-MFN2 (12186-1-AP, Proteintech); anti-AFG3L2 (14631-1-AP, Proteintech); anti-GAPDH (M20028, Abmart); anti-LonP1 (28020S, Cell Signaling Technology); anti-HSP60 (12165, Cell Signaling Technology); anti-PDI (3501P, Cell Signaling Technology); anti-IRE1α (3294P, Cell Signaling Technology); anti-eIF2α (5324S, Cell Signaling Technology); anti-p-eIF2α (3597S, Cell Signaling Technology); anti-ATF4 (11815s, Cell Signaling Technology); anti-ATF6 (65880, Cell Signaling Technology); anti-PCK2 (6924, Cell Signaling Technology); anti-VDAC1 (4661S, Cell Signaling Technology); anti-Drp1 (611113, BD-Pharmingen); anti-OPA1 (612607, BD-Pharmingen); anti-Tid-1 L/S (sc-18820, Santa Cruz); anti-OMA1 (sc-515788, Santa Cruz); anti-SDHA (ab14715, Abcam); anti-Cox I (ab14705, Abcam); anti-Cox IV (ab1460643, Abcam); anti-ClpX (ab168338, Abcam); and anti-FGF21 (ab171941, Abcam). Streptavidin-biotin-peroxidase complex (SABC) and 3,3′-diaminobenzidine tetrahydrochloride (DAB) were purchased from Boster.

Protease (Complete Mini, Cat. No. 11 836 145 001), and phosphatase (PhosphoSTOP, Cat. No. 04 906 837 001)) inhibitor cocktail tablets were purchased from Roche Applied Science. MitoTracker® Green FM (9074) was purchased from Cell Signaling Technology. DAPI (Cat. No. P-36935) and MitoTracker® Red CMXRos (M7512) were obtained from Thermo Fisher Scientific. The Sirius Red staining kit (ab150681) was purchased from Abcam.

### Cell Lines and Cell Culture

The human low-passage HEK293T cell line was obtained from the Cell Bank of the Chinese Academy of Sciences. The H9c2 rat embryonic cardiomyocyte cell line was obtained from the Model Animal Research Center of Nanjing University. HEK293T and H9c2 cells were cultured in Dulbecco’s Modified Eagle’s Medium (DMEM, Life Technologies) supplemented with 10% fetal bovine serum (FBS, Life Technologies) and antibiotics (100 units/ ml penicillin and 100 μg/ml streptomycin) at 37°C in a humidified incubator with 5% CO_2_. LonP1 stable knockdown and control H9c2 cells were cultured in DMEM medium supplemented with 10% FBS, penicillin, streptomycin and puromycin (2.0 μg/ml) in a humidified incubator with 5% CO_2_. Both cell lines were confirmed as mycoplasma free and authenticated by the Cell Bank of the Chinese Academy of Sciences and the Model Animal Research Center of Nanjing University before sending to our lab. Cell lines were also routinely tested and confirmed to be mycoplasma free during this study.

### Hematoxylin and Eosin Staining (H&E)

Heart tissues were fixed, embedded in paraffin and sectioned following standard protocols. The sections were stained with hematoxylin for 15 min and washed in running tap water for 20 min. The sections were then counterstained with eosin from 30 sec to 1 min. Finally, the sections were dehydrated in 95% and absolute ethanol and mounted in Permount (Thermo Fisher Scientific).

### Sirius Red Staining

Sirius Red staining was performed using a Picro Sirius Red staining kit according to the manufacturer’s instructions (Thermo Fisher Scientific). Briefly, paraffin-embedded heart sections were deparaffinized/hydrated, and a series of washes was performed, followed by staining with Picro’s Sirius Red for 1□hour and washing for 1□min in 0.1□N HCl. Slides were dehydrated and mounted in synthetic resin.

### Echocardiography

An experienced researcher blinded to the study performed echocardiographic evaluations to avoid biases. A Vevo 3100 high-resolution micro-ultrasound system (FUJIFILM Visual Sonics Inc.) was used to determine heart function and ventricular dimensions. For this procedure, 1.5% isoflurane was used to anesthetize mice, and then the mice were placed on a heating table in a supine position. M-mode and two-dimensional (2D) images were recorded along a short-axis view from the mid-left ventricle at the tips of the papillary muscles. The interventricular septal thickness (IVS) and left ventricle (LV) internal diameter (LVID) were measured at end-diastole and end-systole. The fractional shortening (FS) and ejection fraction (EF) were calculated from the LV dimensions in the 2D short-axis view.

### Transmission Electron Microscopy (TEM)

The heart tissue samples were fixed with 2.5% (v/v) glutaraldehyde buffer for 15 min at room temperature, followed by overnight at 4°C. The following day, the samples were treated with 1% osmium tetroxide and 0.1 M cacodylate buffer for 1 hour. Samples were stained in 1% uranyl acetate and dehydrated with ethanol. Epoxy-resin-embedded samples were sectioned, and placed on formvar/carbon-coated copper grids. Grids were stained with uranyl acetate and lead nitrate. Then the samples were examined with a JEM1400 transmission electron microscope (JEOL).

### Isolation and Purification of Mitochondria

Heart tissues were separated and washed in Buffer A (0.22 M mannitol, 0.075 M sucrose, 30 mM Tris-HCl, pH 7.4) and then homogenized in Buffer B (0.22 M mannitol, 0.075 M sucrose, 30 mM Tris-HCl, 75 mM BSA, 0.5 mM EGTA, pH 7.4) by 50-100 strokes in a tight-fitting Dounce homogenizer [61]. The lysate was centrifuged at 750 × g for 10 min at 4°C. Next, the supernatant was centrifuged at 9000 × g for 15 min at 4°C, and the precipitate was washed twice with Buffer C (0.22 M mannitol, 0.075 M sucrose, 30 mM Tris-HCl, 75 mM BSA, pH 7.4) and then centrifuged at 10,000 × g for 15 min. The precipitate in the tube was the crude mitochondria, and this mitochondrial fraction was suspended in 200 μl Buffer D (10 mM Tris-HCl, pH 7.4, 1 mM EDTA, 0.32 M sucrose) and 2 μl 100 mM PMSF.

### Blue Native PAGE Gel (BNG)

BNG was performed using a previously published protocol with modifications (39). Briefly, buffer (50 mM NaCl, 50 mM imidazole, 2 mM 6-aminohexanoic, and 1 mM EDTA, pH 7.0) was added to 400 μg of pelleted mitochondria. Then, 12 μl digitonin was added [20% (w/v)], and the mitochondria were solubilized for 10-20 min. After centrifugation at 20,000 × g for 45 min, the supernatants were collected. Next, we added a mixture of glycerol/Coomassie blue G-250 dye (2:1) to the samples to yield a sample/mixture ratio of 5:1 (v/v). Finally, the samples were run on 3-11% acrylamide gradient gels at 4°C.

### Western Blot Analysis

Tissue samples were washed 3 times with ice-cold PBS and homogenized using a homogenizer (Kinematica AG) in 1.5 ml tissue RIPA lysis buffer (50 mM Tris-HCl, pH 7.4, 1.0% Triton X-100, 1% sodium deoxycholate, 0.1% SDS, 150 mM NaCl) supplemented with a protease inhibitor cocktail tablet, and PhosSTOP TM phosphatase inhibitor cocktail tablet. Tissue homogenates were cleared by centrifugation at 18,000 × g for 25 min at 4°C, and the supernatants were collected in clean microcentrifuge tubes on ice. A similar procedure was used to prepare whole-cell extracts from cells. Briefly, cells were washed with ice-cold PBS and lysed in RIPA lysis buffer supplemented with protease and phosphatase inhibitors on ice for 20 min, followed by centrifugation at 18,000 × g for 30 min at 4°C, and the supernatants were collected. Protein concentrations of the tissue homogenates or whole-cell extracts were determined using the Pierce BCA protein assay kit (Thermo Fisher Scientific).

Tissue or cell extracts equivalent to 20 μg total protein were resolved in 10% SDS-PAGE gels followed by electrophoretic transfer onto a nitrocellulose membrane (Bio-Rad) in Tris-glycine buffer. Blots were blocked at room temperature for 1.5 hours in 5% nonfat milk in Tris-buffered saline (TBS)-Tween (TBST) on a shaker, and then incubated with the primary antibodies as indicated in 5% nonfat milk TBST overnight at 4°C. The membrane was washed in TBST at least 3 times for 10 min and then incubated with horseradish peroxidase (HRP)-conjugated anti-rabbit or anti-mouse immunoglobulin G at room temperature for 1 hour with gentle shaking. Immunoreactive proteins were detected by ECL reagent according to the manufacturer’s protocol (Thermo Fisher Scientific). The optical density of the Western blot signals was quantified using the National Institutes of Health (USA) ImageJ software.

### RNA Preparation and Quantitative Real-Time PCR (qRT-PCR)

The total RNA was extracted from heart tissues using TRIzol (Life Technologies) according to the manufacturer’s instructions. The total RNA (1 μg) from each ample was used to synthesize first-strand cDNA by reverse transcription using the PrimeScript™ RT reagent Kit with gDNA Eraser (Takara), according to the manufacturer’s instructions. The cDNA produced was subsequently used as a template for qRT-PCR. qRT-PCR analysis, which was performed using a 2 μl cDNA /20 μl reaction volume on a CFX Connect™ real-time PCR detection system (Bio-Rad) using SYBR Green according to the manufacturer’s protocol. The primer sequences used for qRT-PCR are listed in Supplemental Table 2. Thermal cycling was performed using the following parameters: 95°C for 10 min, then 45 cycles of denaturation at 95°C for 10 sec and extension at 60°C for 30 sec. The threshold cycle number (CT) was recorded for each reaction. The CT value was normalized to that of GAPDH. Each sample was assayed in triplicate and repeated at least three times.

### RNA Interference

psi-LVRU6P vectors (carrying a puromycin antibiotic resistance gene) containing control and shRNA oligonucleotides were purchased from Gene Copoeia (Rockville, MD, USA). The two shRNAs sequences are listed in Supplemental Table 3.

### Lentivirus Production and Transduction

Lentiviral production and transduction were conducted according to the manufacturer’s instructions (Gene Copoeia). Briefly, lentiviral vectors and LonP1 or control shRNA were packed with the Lenti-Pac™ HIV Expression Packaging Kit using HEK293T cells and incubated overnight, followed by replacement of the old culture medium with fresh DMEM supplemented with 5% heat-inactivated FBS and penicillin-streptomycin. Titer Boost reagent (1/500 volume) was added to the culture medium and incubated at 37°C in a humidified incubator with 5% CO2. At 48 hours post transfection, the supernatants containing lentivirus particles were collected, filtered through 0.45 μm syringe filters (Millipore), and used immediately to infect H9c2 cells. To select stably transfected cells, the old medium was replaced by fresh complete medium containing puromycin (2.0 μg/ml) every 3 days until drug-resistant colonies become visible. The knockdown of LonP1 was validated by qRT-PCR and Western blot analysis. Positive clones with stable knockdown of LonP1 were expanded and maintained in medium supplemented with 2.0 μg/ml puromycin.

### Fluorescence Detection in Cultured Cells

For measuring the colocalization of LonP1 with mitochondria, cells were stained with 50 nM MitoTracker Green for 30 min at 37°C and then treated with 4% paraformaldehyde for 5 min, followed by incubation with mouse anti-LonP1 antibody at 4°C overnight. Slides were then incubated with Alexa Fluor 555– conjugated donkey anti-rabbit antibodies (A-31572, Invitrogen) at room temperature for 2 hours. Cells were then stained with DAPI for 30 min at room temperature. Finally, fixed or living cells were visualized by confocal microscopy with a Leica TCS SP8 microscope with a 633 numerical aperture [NA] 1.35 oil objective. To determine mitochondrial morphology, 100 cells were randomly selected for quantitative analysis and visually scored into four classifications (tubular, short tubular, fragmented, and large spherical).

### Cell Proliferation Assay

Control and LonP1 knockdown H9c2 cells were seeded into a 96-well plate at a density of 2 × 103 cells per well and incubated overnight. The viability of H9c2 cells was determined at 1, 2, 3, 4, and 5 days using an MTT Cell Proliferation and Cytotoxicity Assay Kit according to the manufacturer’s protocol.

### XF Extracellular Flux Analyzer Experiments

The intact cellular OCR of LonP1 stable knockdown and control H9c2 cells was measured using a Seahorse XF-96 Extracellular Flux Analyzer (Seahorse Bioscience) as described previously (40). Results were obtained by performing three independent experiments with 8 replicates of LonP1 stable knockdown and control H9c2 cells. After the assay was completed, the protein concentration was determined by a BCA protein assay kit to normalize the OCR according to the manufacturer’s instructions.

### FACS Analysis for ROS and Mitochondrial Membrane Potential

Intracellular ROS levels were measured using the fluorescence probe 2’,7’-dichlorodihydrofluorescein diacetate (DCFH-DA) according to the manufacturer’s protocol (Beyotime). DCFH-DA diffuses into cells and is deacetylated by esterases to nonfluorescent 2’,7’-dichlorofluorescin (DCFH), which is trapped inside the cells and rapidly oxidized by ROS (including H2O2) to form highly fluorescent 2’,7’-dichlorofluorescein (DCF). The fluorescence intensity at 530 nm is proportional to the ROS levels within the cell cytosol. Briefly, Lon stable knockdown H9c2 or vector control cells were trypsinized, washed with DMEM, and incubated with DCFH-DA at a final concentration of 10 μM in DMEM for 30 min at 37°C, then the cells were washed three times with DMEM. Mitochondrial superoxide levels were detected by MitoSOX staining according to the manufacturer’s protocol (Thermo Fisher Scientific). Briefly, control and LonP1 stable knockdown H9c2 cells were incubated with 5 μM MitoSOX Red for 10 min at 37°C and then subjected to flow cytometry analysis. Data are plotted as median fluorescence intensities (MFI). The mitochondrial membrane potential was analyzed by JC-1 staining according to the manufacturer’s protocol (Beyotime). Briefly, LonP1 stable knockdown H9c2 or vector control cells were stained with 2.0 μM JC-1 in complete medium and incubated for 20 min at 37°C in the dark. To remove excess JC-1, cells were washed once with prewarmed PBS and pelleted by centrifugation. Cell pellets were resuspended by gently flicking the tubes, and 500 μl PBS were added to each tube. Cell samples were analyzed immediately using a BD Accuri^TM^ C6 Plus flow cytometer (BD Biosciences). All experiments were carried out at least 3 times independently, with 3 technical replicates in each experiment.

### Statistical Analysis

Statistical analyses were performed with Prism software (GraphPad Prism 5). Data were analyzed using a Student’s t test. P value less than 0.05 was considered as significant. Data with statistical significance (*P < 0.05, **P < 0.01, ***P< 0.001) are shown in the figures. All values are presented as the means ± SEM, obtained from at least three independent experiments.

## Supporting information

Supplemental Experimental Procedures

## Acknowledgements

We thank Dr. Charles Reichman (Rutgers-New Jersey Medical School), and Dr. Jianhong Zhu (Wenzhou Medical University) for their comments on the manuscript and members of the B. L., Z.Y. and G.Y. Lab for technical support and valuable discussions. This study was supported by grants from National Natural Science Foundation of China (31070710, 31171345, 31570772, and 31771534 to B.L., 81774022 to L.J.), and Key Discipline of Zhejiang Province in Medical Technology (First Class, Category A). National Basic Research Program of China (973 Program, 2013CB531702 to B.L., 2013CB531704 to G.Y.)

## Author Contributions

**Conceptualization:** Bin Lu, Zhongzhou Yang, Guanlin Yang

**Data Curation:** Bin Lu, Fugeng Shangguan, Dawei Huang, Shiwei Gong

**Formal analysis:** Bin Lu, Fugeng Shangguan, Dawei Huang, Shiwei Gong, Yingchao Shi, Zhiying Song, Chaojun Yan, Yujie Li, Juan Xu, Shengnan Han, Pingyi Chen, Lu Wang, Carolyn K. Suzuki

**Funding acquisition:** Bin Lu, Lianqun Jia, Guanlin Yang

**Investigation:** Bin Lu, Fugeng Shangguan, Dawei Huang, Shiwei Gong, Yingchao Shi, Yujie Li, Juan Xu, Chaojun Yan, Mingjie Xu, Shengnan Han, Pingyi Chen, Lu Wang

**Methodology:** Dawei Huang, Shiwei Gong, Yingchao Shi, Zhiying Song, Lianqun Jia, Juan Xu, Tongke Chen, Mingjie Xu, Yujie Li, Nan Song, Yongzhang Liu, Xingxu Huang, Zhongzhou Yang, **Project administration:** Bin Lu

**Resource:** Lianqun Jia, Juan Xu, Tongke Chen, Xingxu Huang, Carolyn K. Suzuki, Zhongzhou Yang, Guanlin Yang

**Supervision:** Bin Lu, Zhongzhou Yang, Zhiying Song, Guanlin Yang

**Validation:** Bin Lu, Fugeng Shangguan, Pingyi Chen, Lu Wang

**Visualization:** Bin Lu, Dawei Huang, Shiwei Gong, Zhiying Song, Chaojun Yan, Yujie Li, Yongzhang Liu **Writing – original draft:** Bin Lu, Zhongzhou Yang, Fugeng Shangguan, Dawei Huang, Shiwei Gong, Zhiying Song, Yujie Li

**Writing – review & editing:** Bin Lu, Zhongzhou Yang, Carolyn K. Suzuki, Fugeng Shangguan, Dawei Huang, Shiwei Gong, Yujie Li, Zhiying Song, Pingyi Chen, Xingxu Huang

