## Supplemental Figures for "LonP1 Orchestrates UPR^mt^ and UPR^ER^ and Mitochondrial Dynamics to Regulate Heart Function"

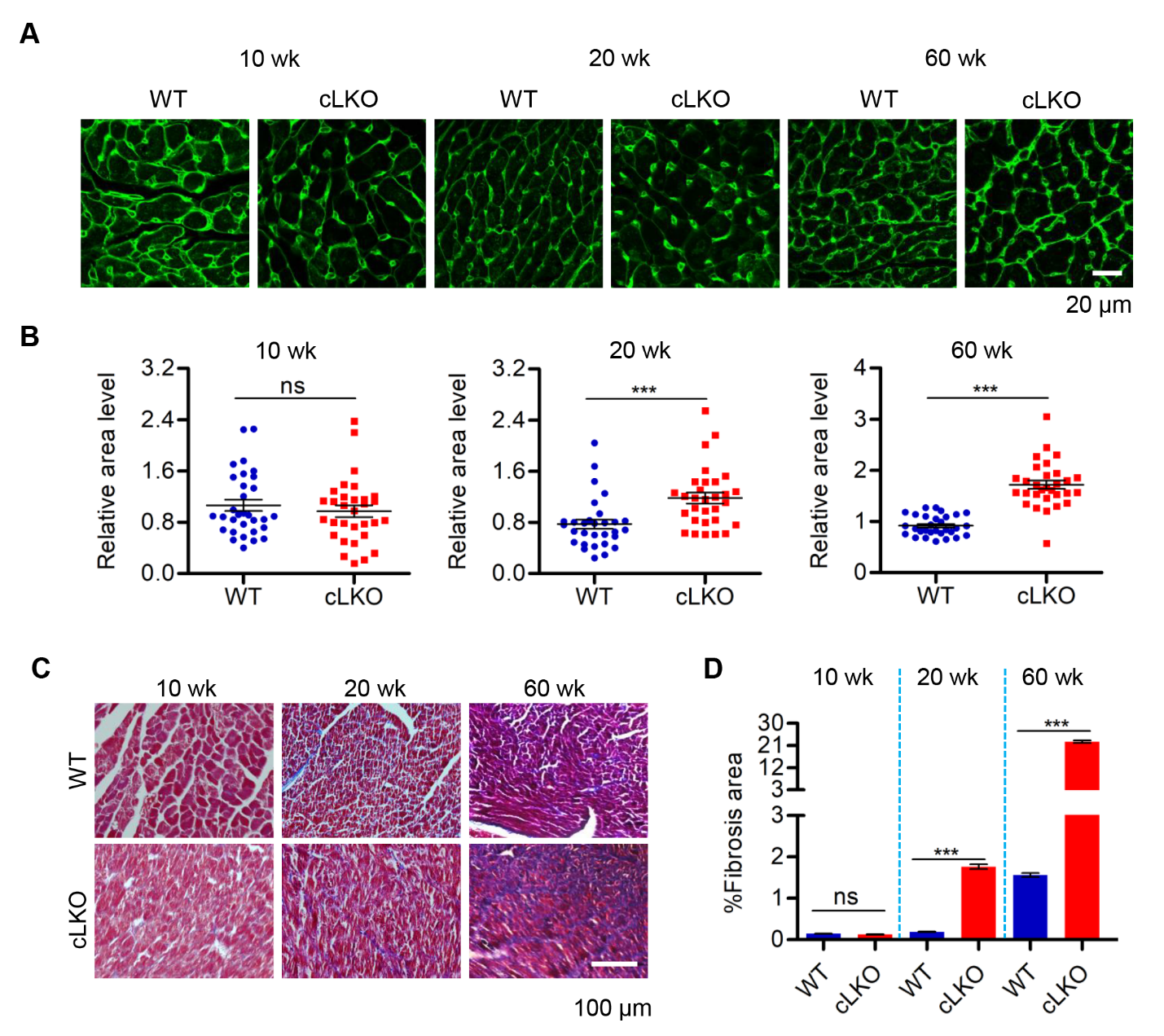

**Figure S1. Cardiomyocyte-Specific Deletion of LonP1 Causes Pathological Heart Remodeling and Myocardial Fibrosis, Related to Figure 1.**

(**A** and **B**) Wheat germ agglutinin stain to outline the cell surface area in myocardial cell from 10-, 20-, and 60-week old WT and cLKO mice. Scale bars, 20 μm. Data are presented as mean ± SEM (n=30, ns, no significant, ****P* < 0.001, statistically significant by Student’s t test).

(**C**) Representative images show Masson’s trichrome staining of the cardiac tissues from 10-, 20- and 60- week-old WT and cLKO mice. Scale bars, 100 μm.

(**D**) Quantitative analyses of the heart fibrotic area in 10-, 20-, and 60-week-old WT and cLKO mice. Data are presented as mean ± SEM (n=3, ns, no significant, ****P* < 0.001, statistically significant by Student’s t test).

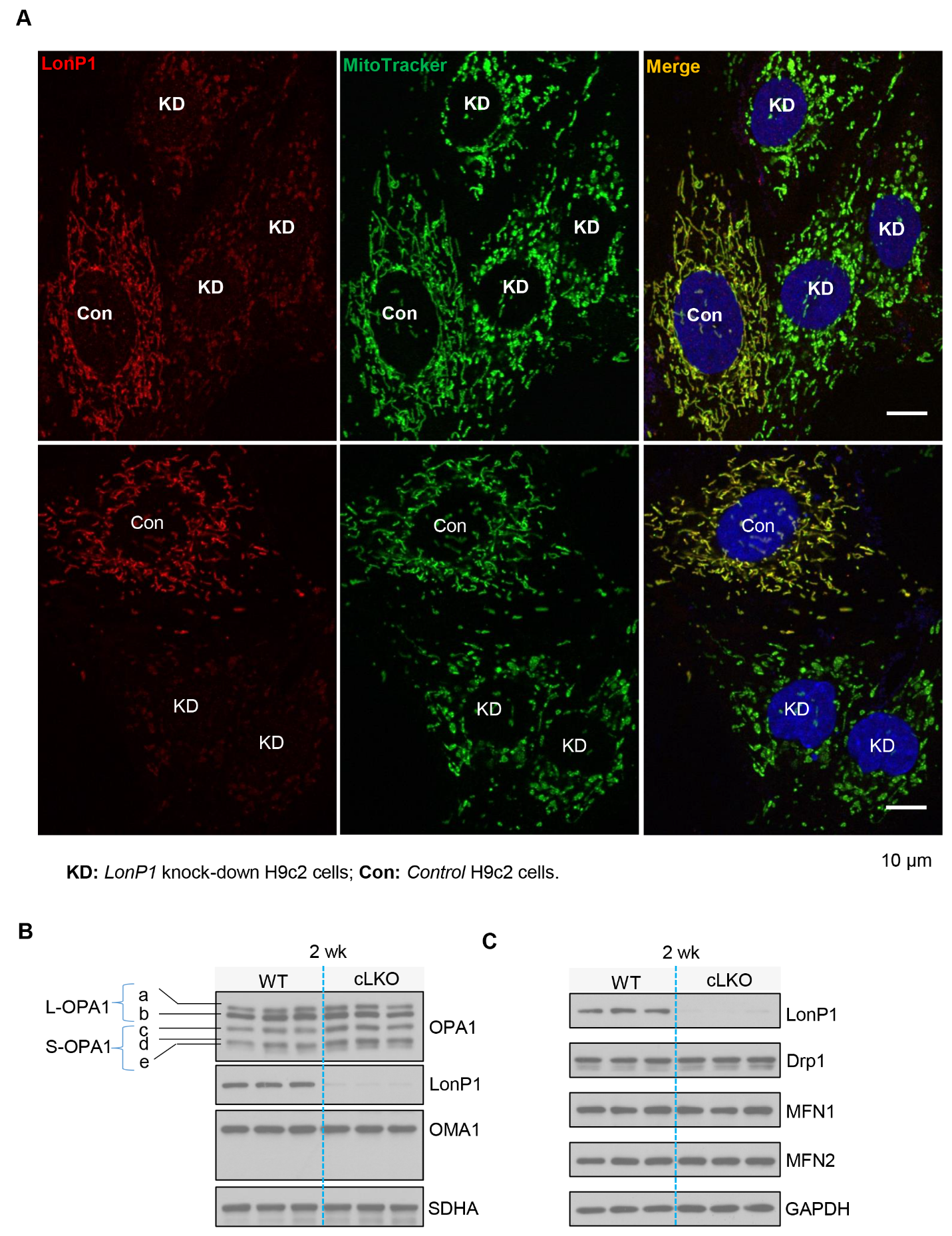

(legend on next page)

**Figure S2. LonP1 Deletion Leads to Mitochondrial Fragmentation, Related to Figure 2.**

(A) Representative confocal microscopy images (scale bars, 10 μm) of mitochondria in sh*Cont* and sh*LonP1* H9c2 cells. sh*Cont* or sh*LonP1* H9c2 cells were immunostained with specific LonP1 antibodies and MitoTracker Green. Colocalization of LonP1 and mitochondria was visualized by confocal microscopy. (B and C) Western blot analysis of mitochondrial fusion- and fission-related proteins OPA1, OMA1, SDHA, DRP1, MFN1, and MFN2, as well as LonP1 protein levels in the heart tissue of WT and cLKO mice at 2 weeks of age. GAPDH was used as a loading control.

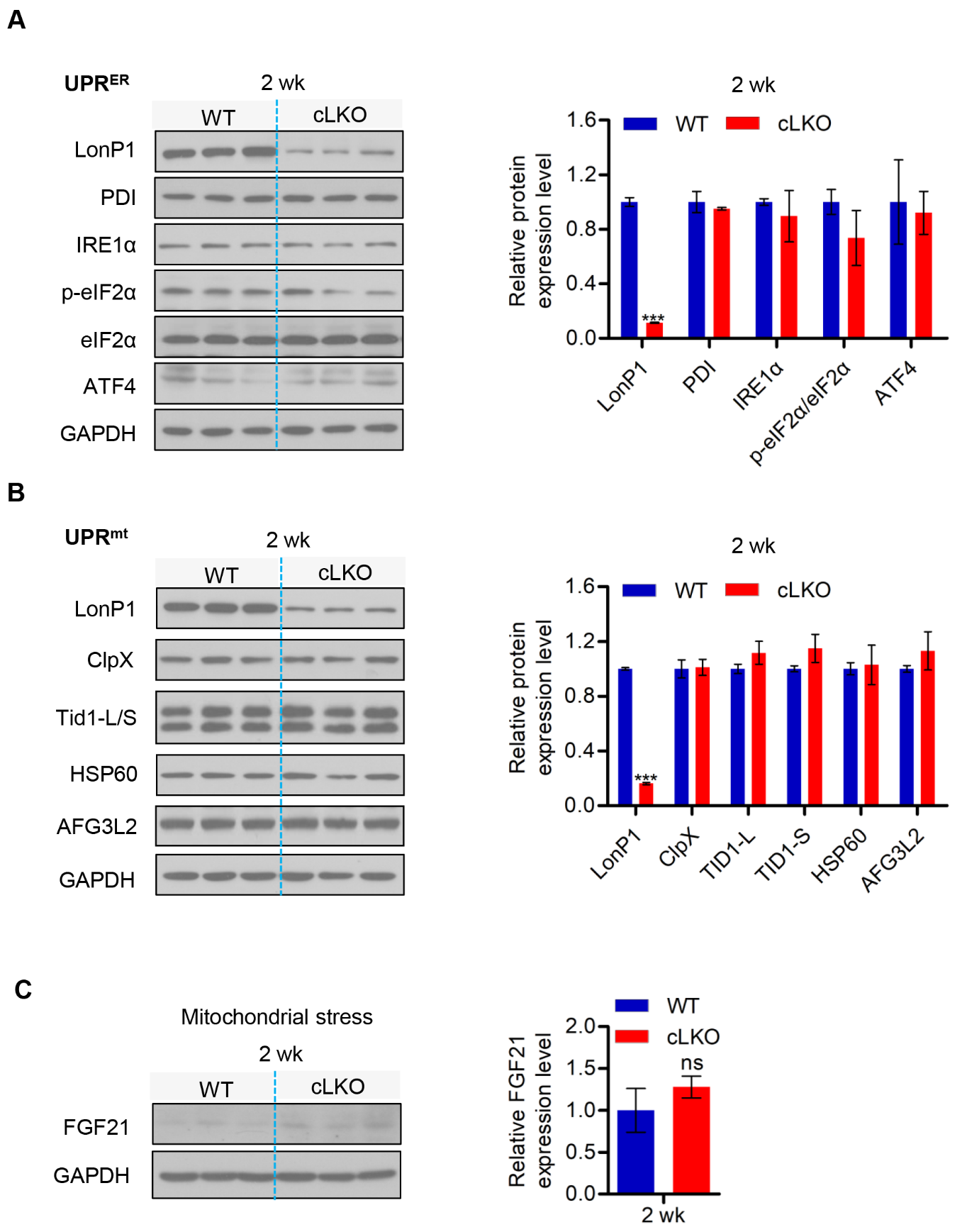

(legend on next page)

**Figure S3. UPR^ER^- and UPR^mt^-Related Proteins, as well as FGF21 Expression Remained Unchanged in the Hearts of WT and cLKO Mice of 2-Week-Old, Related to Figure 3.**

(A) Western blot analysis and quantification of UPR^ER^-related protein levels of PDI, IRE1α, p-eIF2α, ATF4, and LonP1 in the hearts of 2-week-old WT and cLKO mice. GAPDH was used as a loading control.

(B) Western blot analysis and quantification of UPR^mt^-related protein levels of ClpX, Tid1-L/S, HSP60, FG3L2, and LonP1 in the hearts of 2-week-old WT and cLKO mice. GAPDH was used as a loading control.

(C) Western blot analysis and quantification of FGF21 level in the hearts of 2-week-old WT and cLKO mice. GAPDH was used as a loading control. In (A), (B) and (C) , data are presented as the means ± SEM (n = 3, ns, no significant, ****P* < 0.001, statistically significant by Student’s t test).
