## Supplemental Tables for "LonP1 Orchestrates UPR^mt^ and UPR^ER^ and Mitochondrial Dynamics to Regulate Heart Function"

**Table S1. Genotyping Primers of Mice, Related to Figure 1.**

| **Gene** | **Forward primer** | **Reverse primer** |
| --- | --- | --- |
| ***Lonp1^LoxP/LoxP^*** | 5’-AGGATCACCCTGAGTTCCCAGTT-3’ | 5’-CACCACCTATAGCAGGTGCGAA-3’ |
| ***α-MHC-cre*** | 5’-GCCTGCATTACCGGTCGATGC-3’ | 5’-CAGGGTGTTATAAGCAATCCC-3’ |

**Table S2. Primer Sequence for qRT-PCR, Related to Figure 5.**

| **Gene** | | **Forward** | | **Reverse** |
| --- | --- | --- | --- | --- |
| ***GLS*** | 5’-TTCCAGAAGGCACAGACATGGTTG-3’ | | 5’-GCCAGTGTCGCAGCCATCAC-3’ | |
| ***Suclg2*** | 5’-CAGCGAACTTCTTGGACCTTGGAG-3’ | | 5’-TCCGTTGGCAATGATGGCACAG-3’ | |
| ***Pck2*** | 5’-GCGGCTATGGTGGTAACTCCTTG-3’ | | 5’-GCCAGATTGGTCTTGCCACAGG-3’ | |
| ***Phgdh*** | 5’-CAGGTGGTGGAGAAGCAGAACTTG-3’ | | 5’-GCAGCCTCCAGATCCACATTGTC-3’ | |
| ***Psat1*** | 5’-TCGCTGGTGCTCAGAAGAATGTTG-3’ | | 5’-CTTGATCCATTCCAGGACCATGCC-3’ | |

**Table S3. shRNA sequences of *LonP1*.**

| ***shRNA*** | | | **Forward** | | **Reverse** |
| --- | --- | --- | --- | --- | --- |
|  | ***shLonP1* #1** | 5’-GATCCGGCGCTTTATCAAGATCGT  GGATCAAGAGCGCGAAATAGTTCTA  GCACGTTTTTTTGGAATT-3’ | | 5’-CTAGGCCGCGAAATAGTTCTAGCA  CGTAGTTCTCGCGCTTTATCAAGATCGTGGAAAAAAACCTTAA-3’ | |
|  | ***shLonP1* #2** | 5’-GATCCGCGTTCGCTCAGATTCATG  AGATCAAGAGGCAAGCGAGTCTAAG  TACTCTTTTTTTGGAATT-3’ | | 5’-CTAGGCGCAAGCGAGTCTAAGTA  CTCTAGTTCTCCGTTCGCTCAGATT  CATGAGAAAAAAACCTTAA-3’ | |
