## Supplemental Experimental Procedures for "LonP1 Orchestrates UPR^mt^ and UPR^ER^ and Mitochondrial Dynamics to Regulate Heart Function"

**Reagents and Antibodies**

BCA Protein Assay Kit and Pierce ECL Western Blotting Substrate were obtained from Thermo Fisher Scintific. Primary antibodies included anti-MFN1 (13798-1-AP, Proteintech); anti-MFN2 (12186-1-AP, Proteintech); anti-AFG3L2 (14631-1-AP, Proteintech); anti-GAPDH (M20028, Abmart); anti-LonP1 (28020S, Cell Signaling Technology); anti-HSP60 (12165, Cell Signaling Technology); anti-PDI (3501P, Cell Signaling Technology); anti-IRE1α (3294P, Cell Signaling Technology); anti-eIF2α (5324S, Cell Signaling Technology); anti-p-eIF2 α (3597S, Cell Signaling Technology); anti-ATF4 (11815s, Cell Signaling Technology); anti-Drp1 (611113, BD-Pharmingen); anti-OPA1 (612607, BD-Pharmingen); anti-Tid-1 L/S (sc-18820, Santa Cruz); anti-OMA1 (sc-515788, Santa Cruz); anti-SDHA (ab14715, Abcam); anti-ClpX (ab168338, Abcam); and anti-FGF21 (ab171941, Abcam). Protease (Complete Mini) and phosphatase (PhosphoSTOP^TM^) inhibitor cocktail tablets were purchased from Roche Applied Science. MitoTracker® Green FM (9074) was purchased from Cell Signaling Technology, and MitoTracker® Red CMXRos (M7512) was obtained from Thermo Fisher Scientific. The trichrome stain kit (ab150686) was purchased from Abcam. Wheat germ agglutinin (W834) was purchased from Thermo Fisher Scientific.

**Wheat Germ Agglutinin (WGA) Staining**

Mouse heart tissues were fixed, embedded and sectioned as aforementioned. After deparaffinization and rehydration, sections were stained with 20 mg/ml FITC conjugated-wheat germ agglutinin at room temperature for 30 min, washed with PBS three times and sealed with 50% glycerin.

**Masson Trichrome Stain (Connective Tissue Stain)**

Masson trichrome staining was performed using a Trichrome Stain Kit (Abcam, ab150686) according to the manufacturer’s instructions. Briefly, paraffin-embedded heart section was deparaffinized/hydrated, and a series of washes was performed. Then, the slides were placed in preheated Bouin's fluid for 60 min, followed by a 10 min cooling period. Equal parts of Weigert’s (A) and Weigert’s (B) were mixed, and slides were stained with working Weigert’s iron hematoxylin for 5 min, then rinsed slide in running tap water for 2 min. Then, Biebrich Scarlet /Acid Fuchsin solution was applied to the slides for 15 min. The slides were rinsed in distilled water and differentiated in phosphomolybdic/phosphotungstic acid solution for 10-15 min or until collagen was not red. Without rinsing, aniline blue solution was applied to the slides for 5-10 min, and the slides were rinsed in distilled water. Then, acetic acid solution (1%) was applied to the slides for 3-5 min. Finally, the slides were dehydrated very quickly in 2 changes of 95% alcohol, followed by 2 changes of absolute alcohol, then cleared in xylene and mounted in synthetic.

**Western Blot Analysis**

Tissue samples were washed 3 times with ice-cold PBS and homogenized using a homogenizer (Kinematica AG) in 1.5 ml tissue RIPA lysis buffer (50 mM Tris-HCl, pH 7.4, 1.0% Triton X-100, 1% sodium deoxycholate, 0.1% SDS, 150 mM NaCl) supplemented with a protease inhibitor cocktail tablet, and PhosSTOP phosphatase inhibitor cocktail tablets. Tissue homogenates were cleared by centrifugation at 18,000 × g for 25 min at 4°C, and the supernatants were collected in clean microcentrifuge tubes on ice. A similar procedure was used to prepare whole-cell extracts from cells. Briefly, cells were washed with ice-cold PBS and lysed in RIPA lysis buffer supplemented with protease and phosphatase inhibitors on ice for 20 min, followed by centrifugation at 18,000 × g for 30 min at 4°C, and the supernatants were collected. Protein concentrations of the tissue homogenates or whole cell extracts were determined using the Pierce BCA protein assay kit. Tissue or cell extracts equivalent to 20 μg total protein were resolved in 10% SDS-PAGE gels followed by electrophoretic transfer onto a nitrocellulose membrane (Bio-Rad) in Tris-glycine buffer. Blots were blocked at room temperature for 1.5 hours in 5% nonfat milk in Tris-buffered saline (TBS)-Tween (TBST) on a shaker and then incubated with the indicated primary antibodies in 5% nonfat milk TBST overnight at 4°C. The membrane was washed in TBST at least 3 times for 10 min and then incubated with horseradish peroxidase (HRP)-conjugated anti-rabbit or anti-mouse immunoglobulin G at room temperature for 1 hour with gentle shaking. Immunoreactive proteins were detected by ECL reagent according to the manufacturer’s protocol (Thermo Fisher Scientific). The optical density of the Western blot signals was quantified using the National Institutes of Health ImageJ software.

**Protein Extraction and Trypsin Digestion**

Samples were first ground in liquid nitrogen, and then the cell powder was transferred to a 5-ml centrifuge tube and sonicated three times on ice using a high-intensity ultrasonic processor (Scientz) in lysis buffer (8 M urea, 1% Triton-100, 65 mM DTT and 0.1% Protease Inhibitor Cocktail). The remaining debris was removed by centrifugation at 20,000 × g at 4 °C for 10 min. Finally, the protein was precipitated with cold 15% TCA for 2 hours at -20 °C. After centrifugation at 4 °C for 10 min, the supernatant was discarded. The remaining precipitate was washed with cold acetone three times. The protein was redissolved in buffer (8 M urea, 100 mM TEAB, pH 8.0), and the protein concentration was determined with a 2D Quant kit according to the manufacturer’s instructions. For digestion, the protein solution was reduced with 10 mM DTT for 1 hour at 37 °C and alkylated with 20 mM IAA for 45 min at room temperature in darkness. For trypsin digestion, the protein sample was diluted by adding 100 mM TEAB to a urea concentration of less than 2 M. Finally, trypsin was added at a 1:50 trypsin-to-protein mass ratio for the first digestion overnight and at a 1:100 trypsin-to-protein mass ratio for a second 4 hour-digestion. Approximately 100 μg protein from each sample was digested with trypsin for the following experiments.

**iTRAQ Labeling and LC-MS/MS Analysis**

After trypsin digestion, peptides were desalted by a Strata X C18 SPE column (Phenomenex) and vacuum-dried. Peptides were reconstituted in 0.5 M TEAB and processed according to the manufacturer’s protocol for the 8-plex iTRAQ kit. Briefly, one unit of iTRAQ reagent (defined as the amount of reagent required to label 100 μg of protein) was thawed and reconstituted in 24 μl ACN. The peptide mixtures were then incubated for 2 hour at room temperature and then pooled, desalted and dried by vacuum centrifugation. The sample was then fractionated by high-pH reverse-phase HPLC using an Agilent 300Extend C18 column (5 μm particles, 4.6 mm ID, 250 mm length). Briefly, peptides were first separated with a gradient of 2% to 60% acetonitrile in 10 mM ammonium bicarbonate pH 10 over 80 min into 80 fractions. Then, the peptides were combined into 18 fractions and dried by vacuum centrifugation. Peptides were dissolved in 0.1% FA and directly loaded onto a reversed-phase precolumn (Acclaim PepMap 100, Thermo Fisher Scientific). Peptide separation was performed using a reversed-phase analytical column (Acclaim PepMap RSLC, Thermo Fisher Scientific). The gradient was comprised of an increase from 7% to 20% solvent B (0.1% FA in 98% ACN) over 22 min, then 20% to 35% over 6 min, followed by climbing to 80% over 3 min and holding at 80% for the last 4 min, all at a constant flow rate of 300 nl/min on an EASY-nLC 1000 UPLC system. The resulting peptides were analyzed by a Q Exactive^TM^ Plus hybrid quadrupole-orbitrap mass spectrometer (Thermo Fisher Scientific). The peptides were subjected to an NSI source followed by tandem mass spectrometry (MS/MS) in Q Exactive^TM^ plus (Thermo Fisher Scientific) coupled online to the UPLC. Intact peptides were detected in the orbitrap at a resolution of 70,000. Peptides were selected for MS/MS using the NCE setting of 33, and ion fragments were detected in the orbitrap at a resolution of 17,500. A data-dependent procedure that alternated between one MS scan followed by 20 MS/MS scans was applied for the top 20 precursor ions above a threshold ion count of 2E4 in the MS survey scan with 10.0s dynamic exclusion. The electrospray voltage applied was 2.0 kV. Automatic gain control (AGC) was used to prevent overfilling of the ion trap, and 5E4 ions were accumulated for the generation of MS/MS spectra. For MS scans, the m/z scan range was 350 to 1800. The fixed first mass was set as 100 m/z. The resulting MS/MS data were processed using the Mascot search engine (v.2.3.0). Tandem mass spectra were searched against the UniProt rat rattus database (33,648 sequences). Trypsin/P was specified as the cleavage enzyme, allowing up to 2 missing cleavages. The mass error was set to 10 ppm for precursor ions and 0.02 Da for fragment ions. Carbamidomethyl on Cys, iTRAQ-8plex (N-term) and iTRAQ-8plex (K) were specified as fixed modifications, and oxidation on Met was specified as the variable modification. The FDR was adjusted to < 1%, and the peptide ion score was set at > 20.

**Statistical Analysis**

Statistical analyses were performed with Prism software (GraphPad Prism 5.0). Data were analyzed using a Student’s t test and One-way analysis of variance (ANOVA). Differences were considered statistically significant when the p-value was less than 0.05. Data with statistical significance (*p < 0.05, **p < 0.01, ***p< 0.001) are shown in Figures. All values are presented as means ± SEM, obtained from at least three independent experiments.
